# Stem cell homeostasis in the root of *Arabidopsis* involves cell type specific complex formation of key transcription factors

**DOI:** 10.1101/2024.04.26.591257

**Authors:** Vivien I. Strotmann, Monica L. García-Gómez, Yvonne Stahl

**Affiliations:** Institute for Developmental Genetics, Heinrich-Heine University, Universitätsstraße 1, 40225 Düsseldorf, Germany; Cluster of Excellence on Plant Sciences (CEPLAS), Heinrich-Heine University, Universitätsstraße 1, 40225 Düsseldorf, Germany; Theoretical Biology and Bioinformatics (IBB), Utrecht University, Padualaan 8, 3584 CS Utrecht, The Netherlands; Experimental and Computational Plant Development (IEB), Utrecht University, Padualaan 8, 3584 CS Utrecht, The Netherlands; CropXR Institute, The Netherlands

**Keywords:** mathematical modelling, root stem cell niche, FRET-FLIM, transcription factor complexes, PLTs, WOX5, BRAVO

## Abstract

In *Arabidopsis thaliana*, the stem cell niche (SCN) within the root apical meristem (RAM) is maintained by an intricate regulatory network that ensures optimal growth and high developmental plasticity. Yet, many aspects of this regulatory network of stem cell quiescence and replenishment are still not fully understood. Here, we investigate the interplay of the key transcription factors (TFs) BRASSINOSTEROID AT VASCULAR AND ORGANIZING CENTRE (BRAVO), PLETHORA 3 (PLT3) and WUSCHEL-RELATED HOMEOBOX 5 (WOX5) involved in SCN maintenance. Phenotypical analysis of mutants involving these TFs uncover their combinatorial regulation of cell fates and divisions in the SCN. Moreover, interaction studies employing fluorescence resonance energy transfer fluorescence lifetime imaging microscopy (FRET-FLIM) in combination with novel analysis methods, allowed us to quantify protein-protein interaction (PPI) affinities as well as higher-order complex formation of these TFs. We integrated our experimental results into a computational model, suggesting that cell type specific profiles of protein complexes and characteristic complex formation, that is also dependent on prion-like domains in PLT3, contribute to the intricate regulation of the SCN. We propose that these unique protein complex ‘signatures’ could serve as a read-out for cell specificity thereby adding another layer to the sophisticated regulatory network that balances stem cell maintenance and replenishment in the *Arabidopsis* root.

## Introduction

As sessile organisms, plants must cope with environmental challenges and adapt their growth and development accordingly, as they cannot escape adverse conditions. The root system of higher plants plays a pivotal role for the plant’s fitness, as it provides anchorage to the soil and access to water and nutrients. To ensure high developmental plasticity, plants maintain a reservoir of stem cells that reside in the root apical meristem (RAM) at the tip of the root. In *Arabidopsis thaliana* (*A. thaliana*), the center of the RAM harbours a group of slowly dividing, pluripotent stem cells termed the quiescent centre (QC). The QC exerts two key functions: first it produces the surrounding tissue-specific stem cells, also referred to as initials, which by asymmetric cell divisions give rise to different cell types from the outside to the inside: epidermis/lateral root cap, cortex, endodermis, pericycle and stele, as well as the columella at the root tip (Fig. 1 G). Second, the QC serves as signalling hub to maintain the surrounding stem cells in a non-cell autonomous manner (Dolan *et al*., 1993; van den Berg *et al*., 1997; Benfey and Scheres, 2000). The balance between QC quiescence and stem cell replenishment has to be maintained throughout the entire life cycle of a plant and therefore requires fine-tuned regulation, necessitating phytohormones, receptors and their ligands as well as several key transcription factors (TFs) (García-Gómez *et al*., 2021; Strotmann and Stahl, 2021).

**Figure 1:**
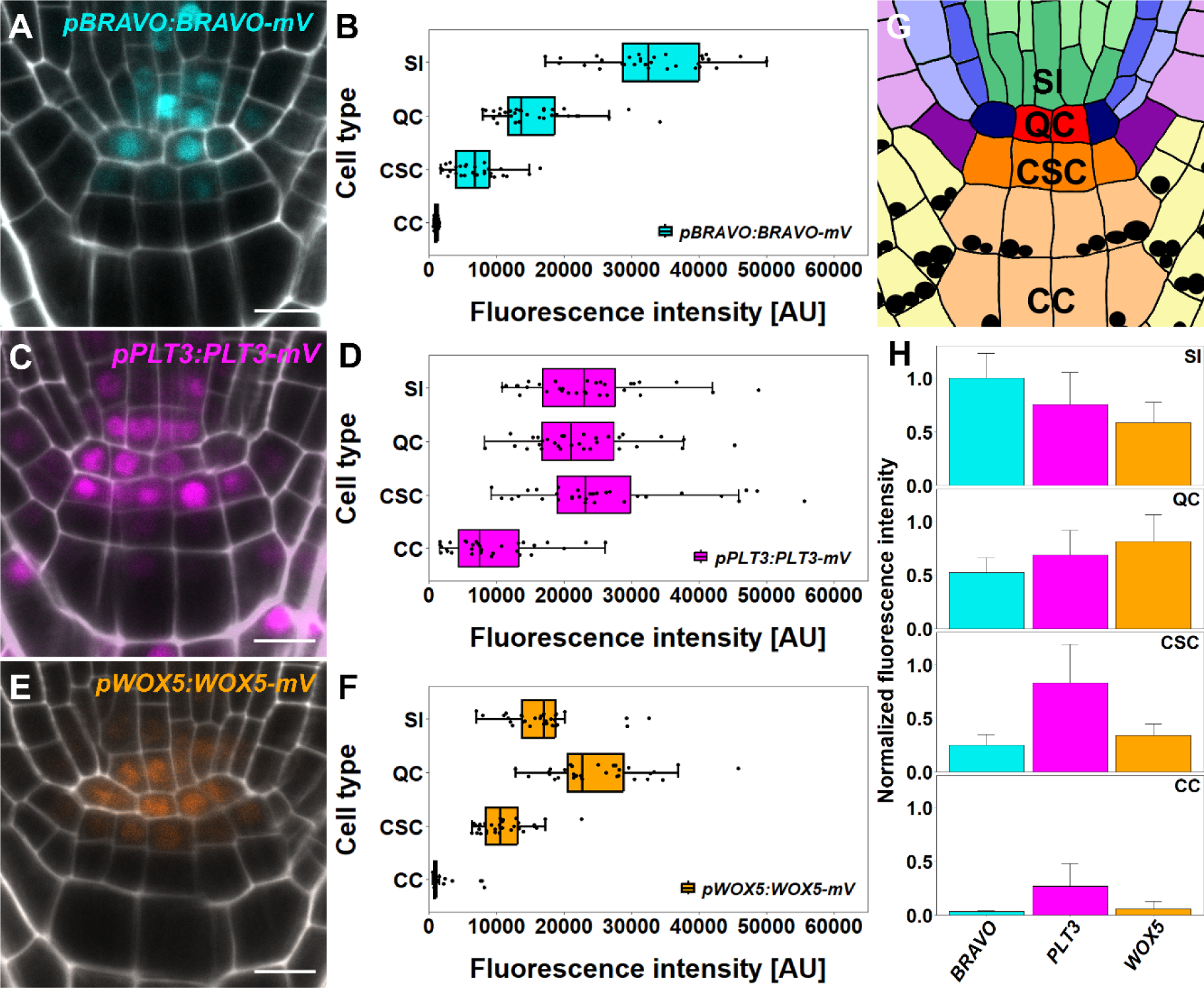
Abundance of BRAVO, PLT3 and WOX5 in the Arabidopsis RAM. Representative images of translational reporter of **A)** BRAVO, **C)** PLT3 and **E)** WOX5 in wildtype *Col-0* background in the RAM as well as the cell type specific quantification of mVenus (mV) fluorescence intensity in **B)** for BRAVO, **D)** for PLT3 and **F)** for WOX5. **G)** Schematic overview of the organisation of the Arabidopsis RAM. The different cell types are represented by different colours. QC: red, cortex endodermis initial: dark blue, endodermis: mid blue, cortex: light blue, stele initials (SI): green, stele: light green, lateral root cap/epidermis initial: purple, epidermis: light purple, lateral root cap: light yellow, columella stem cell (CSC): orange and columella cell (CC): light orange. Starch granules are visualised as black dots. **H)** Bar plot representing the mean fluorescence intensities of mV in BRAVO, PLT3 or WOX5 translational reporters in SIs, QCs, CSCs and CCs normalized to the maximum intensity found for BRAVO in SIs. Error bars display standard deviation. Cell walls were stained using PI and are shown in white, expression of TF is visualized by mVenus in cyan (BRAVO), pink (PLT3) or orange (WOX5). Scalebars represent 10 µm.

The homeodomain TF WUSCHEL-RELATED HOMEOBOX 5 (WOX5) was shown to act as a key regulator for stem cell maintenance in the root (Sarkar *et al*., 2007). By repressing *CYCLIN D3;3* (*CYCD3;3*) and *CYCLIN D1;1* (*CYCD1;1*), WOX5 inhibits periclinal cell divisions in the QC (Forzani *et al*., 2014). Furthermore, WOX5 preserves the undifferentiated status of the columella stem cells (CSCs) by repressing *CYCLING DOF FACTOR 4* (*CDF4*), which involves the recruitment of TOPLESS (TPL) and HISTONE DEACETYLASE 19 (HDA19) (Pi *et al*., 2015). Recent findings suggest that to control the balance between maintaining the stem cell fate of CSCs and their differentiation, WOX5 also interacts with the auxin-dependent APETALA2-type TF PLETHORA 3 (PLT3) (Burkart *et al*., 2022). The *PLT* gene family comprises six members that are described as master regulators of root development (Aida *et al*., 2004; Galinha *et al*., 2007; Mähönen *et al*., 2014). While PLT5 and 7 are mainly involved in lateral root development (Hofhuis *et al*., 2013; Du and Scheres, 2017), PLT1-4 are expressed in the main root forming instructive protein gradients that are necessary for correct QC positioning and cell fate decisions (Aida *et al*., 2004; Galinha *et al*., 2007; Mähönen *et al*., 2014). Interestingly, loss of PLT3 or WOX5 function, as observed in *plt3-1* and *wox5-1* mutants, causes an increase of QC divisions (Sarkar *et al*., 2007; Pi *et al*., 2015; Burkart *et al*., 2022). This phenotype is even more severe in the *plt3 wox5* double mutant indicating that PLT3 and WOX5 act in parallel pathways to control stem cell maintenance in the root (Burkart *et al*., 2022).

In the past decade the brassinosteroids (BRs), a class of phytohormones, were described to play an important role in the regulation of the root stem cell niche (SCN) maintenance (González-García *et al*., 2011). In the *Arabidopsis* RAM, BRs act via the R2R3-MYB TF BRASSINOSTEROIDS AT VASCULAR AND ORGANIZING CENTRE (BRAVO) which inhibits QC divisions and is negatively regulated by the BR-dependent repressor complex formed by BRI1-EMS-SUPPRESSOR 1 (BES1) and TPL on transcript and protein level (Vilarrasa-Blasi *et al*., 2014; Espinosa-Ruiz *et al*., 2017). Recently, the ability of BRAVO to control formative QC divisions was linked to WOX5 (Betegón-Putze *et al*., 2021), as *bravo-2* mutants, like *wox5-1* mutants, show an increased frequency of QC divisions (Sarkar *et al*., 2007; Pi *et al*., 2015; Betegón-Putze *et al*., 2021; Burkart *et al*., 2022).

In addition to the described genetic interactions, one-on-one protein-protein interactions (PPIs) have been reported for WOX5 and PLT3 as well as for BRAVO and WOX5 (Betegón-Putze *et al*., 2021; Burkart *et al*., 2022). However, It is still unknown whether these TFs can also form higher order complexes. Additionally, it remains elusive how these genetic and physical interactions could possibly influence the regulation of stem cell maintenance. To unravel the underlying interplay of key TFs in the root SCN, we used an integrative experimental and computational approach to analyze the protein complex formation between WOX5, PLT3 and BRAVO in the cells of the root SCN. Here, we show that cell type specific profiles of protein complexes are formed and align their occurrence with phenotypical SCN defects of the respective mutants. Moreover, by the deletion of specific interaction sites, we could demonstrate that heterodimerization contributes to maintaining stem cells in the root. Altogether, our results suggest that these unique protein complex ‘signatures’ convey cell type specificity and could explain the different roles played by BRAVO, PLT3 and WOX5 in root SCN maintenance.

## Results

### BRAVO, PLT3 and WOX5 exhibit cell type specific differences in protein abundance in the root SCN

First, we analysed the absolute and relative abundance of BRAVO, PLT3 and WOX5 protein levels in the different cell types found in the SCN of the *Arabidopsis* root, focusing on the stele initials (SIs), QC, CSCs and columella cells (CCs) (Fig. 1 G), by measuring the fluorescence intensity of mVenus (mV) in nuclei of the previously described *pPLT3:PLT3-mV* and *pWOX5:WOX5-mV* translational reporters in *Col-0* WT background (Burkart *et al*., 2022). Additionally, we generated a stable transgenic *Arabidopsis* line expressing *pBRAVO:BRAVO-mV* also in the *Col-0* WT background. We used the same microscopy settings for these quantifications to ensure that the detected protein levels are comparable. Consistent with previous findings, BRAVO protein levels are highest in the SIs and gradually decrease towards the CCs (Fig. 1 A, B) (Vilarrasa-Blasi *et al*., 2014). PLT3 levels are similar in SIs, QC and CSCs, but notably lower in the CCs (Fig. 1 C, D). WOX5 protein levels peak in the QC, decrease in the adjacent SIs and CSCs and are almost completely absent in CCs (Fig. 1 E, F).

We summarized our findings in a protein abundance profile for each individual cell type displaying relative protein levels of BRAVO, PLT3 and WOX5. The protein levels are normalized to the overall maximum intensity, which was found for BRAVO in SIs (Fig. 1 H). Accordingly, We found that BRAVO is the most abundant protein in the SIs, followed by PLT3 and WOX5 in descending order. Conversely, in the QC, we observe a contrasting pattern, marked by WOX5 as the most abundant protein, followed by PLT3 and BRAVO. PLT3 emerges as the predominant protein in the adjacent CSCs, accompanied by low levels of WOX5 and BRAVO protein. In differentiated CCs, WOX5 and BRAVO are almost absent and only low levels of PLT3 can be found. Interestingly, while all of these regulators are expressed in several root SCN cells, our observations reveal quantitative differences in protein abundance that can be combined into a cell type specific ‘fingerprint’. This provides a comprehensive snapshot of the unique protein levels within each cellular context, which could act as an instructive output of cell type specification (Fig. 1 H).

### BRAVO, PLT3 and WOX5 jointly control CSC fate and QC divisions

Several studies have highlighted the inhibitory effect of BRAVO, PLT3 and WOX5 on QC divisions and CSC differentiation in the *Arabidopsis* root (Aida *et al*., 2004; Galinha *et al*., 2007; Vilarrasa-Blasi *et al*., 2014; Mähönen *et al*., 2014; Forzani *et al*., 2014; Pi *et al*., 2015). While all three proteins have been demonstrated to be present in the QC and CSCs, a combinatory effect on QC division and CSC fate has only been demonstrated for WOX5 and PLT3 (Burkart *et al*., 2022) as well as for WOX5 and BRAVO (Betegón-Putze *et al*., 2021). Notably, such interplay has not been observed for BRAVO and PLT3, nor for the simultaneous involvement of all three proteins.

To address this, we have performed SCN stainings, that combines 5-ethynyl-2’- deoxyuridine (EdU) and modified pseudo Schiff base propidium iodide (mPS-PI) stainings (Burkart *et al*., 2022), in several single and multiple mutants. This allowed us to analyse the differentiation status of the distal meristem, as well as the number of QC divisions that occurred within the last 24 h within the same root. To quantify CSC layers, the number of cell layers that lack starch granules distally to the QC were counted. In *Col-0* WT, 68 % of the roots show one CSC layer, whereas only 2 % of the roots lack the starch-free CSC layer and 30 % show two CSC layers, most likely because they have recently divided (Fig. 2 A, B, J, Fig. S1 A). In *bravo-2* and *plt3-1* single mutants, the number of roots showing no CSC layer increases to 11 % and 12 %, respectively (Fig. 2 C, D, J, Fig. S1 B, C). Interestingly, the number of roots displaying no starch-free CSC layer increased to 37 % in *bravo plt3* double mutants (Fig. 2 F, J, Fig. S1 F). This additive effect indicates that PLT3 and BRAVO act in parallel pathways to control CSC differentiation. In 53 % of the *wox5-1* mutant roots, the starch-free CSC layer is missing (Fig. 2 E, J, Fig. S1 D), further emphasizing the importance of WOX5 for CSC fate (Sarkar *et al*., 2007; Pi *et al*., 2015; Burkart *et al*., 2022). Additionally, the *bravo wox5* and the *plt3 wox5* double mutants show an even higher percentage of roots lacking the starch-free CSC, 90 % and 74 % respectively, compared to the single mutants and the *bravo plt3* double mutants (Fig. 2 G, H, J, Fig. S1 E, G). On the other hand, the *bravo plt3 wox5* triple mutant, with 85 % of the roots having a differentiated CSC layer, resembles the *bravo wox5* and *plt3 wox5* double mutants (Fig. 2 I, J, Fig. S1 H). These results suggest that BRAVO, PLT3 and WOX5 jointly control CSC fate.

**Figure 2:**
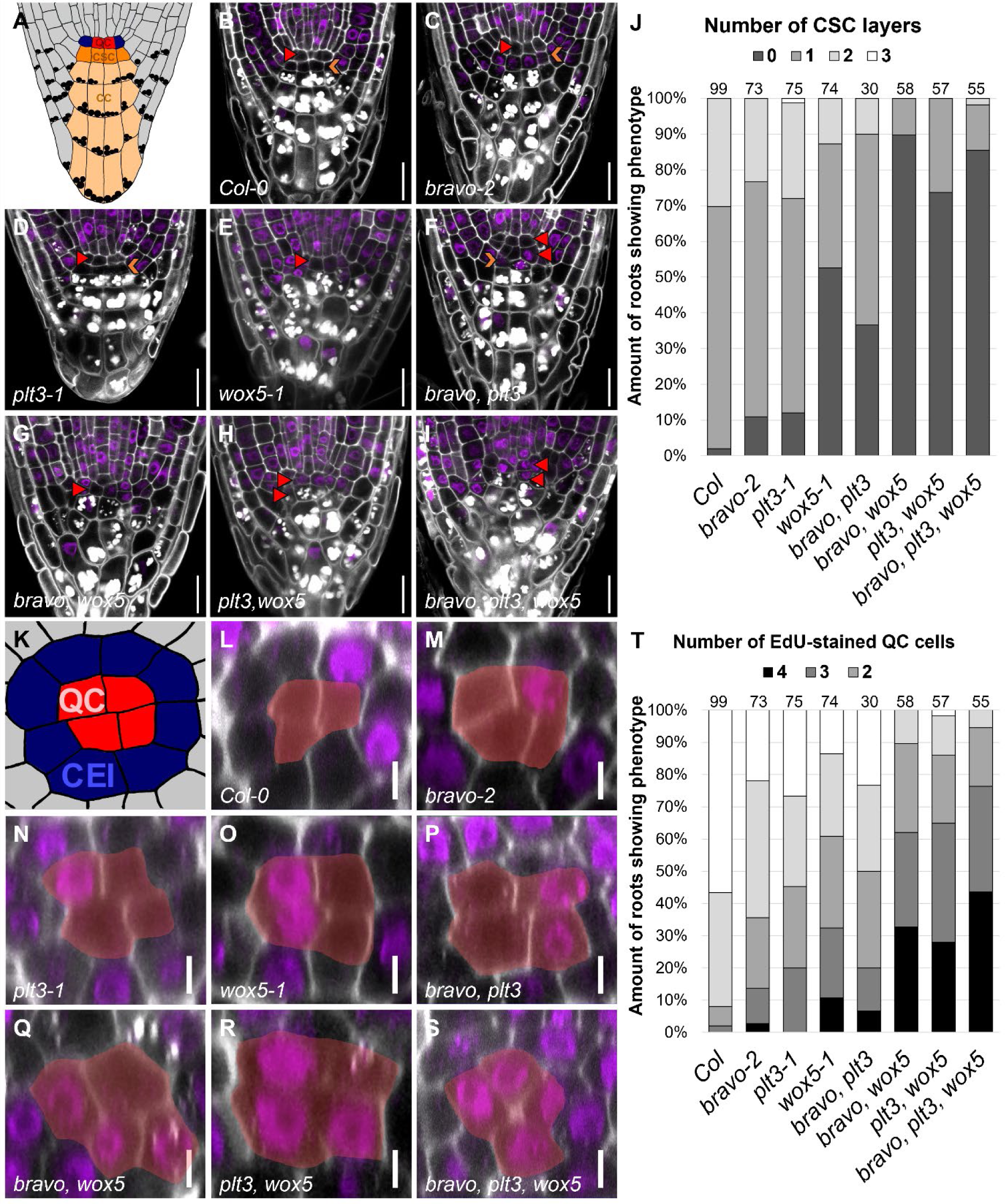
BRAVO, PLT3 and WOX5 jointly regulate CSC differentiation and QC quiescence. A) Schematic representation of a longitudinal section of the Arabidopsis RAM. Red: QC, blue: CEI, dark orange: CSC, light orange: CC. **B-I)** Representative images of the mutant CSC phenotype in the indicated mutant background after combined mPSPI (white) EdU (purple) staining. The position of the QC is indicated by a red arrowhead and the CSC layer is marked with an orange arrowhead. Scale bars represent 20 µm. **J)** Quantification of SCN staining displaying 0, 1, 2 or 3 layers of CSC. The number of analyzed roots for each genotype is indicated above each bar and results from to 3-5 technical replicates. **K)** Schematic representation of a transversal section of the Arabidopsis RAM. QC cells are highlighted in red and CEIs are displayed in blue. **L-S)** Representative images of optical cross sections of the Arabidopsis RAM in the indicated mutant background. The combined mPSPI/EdU staining reveals the cells that have divided within 24 h. QC is highlighted in yellow. Scale bars represent 5 µm **T)** Quantification of SCN staining displaying 0, 1, 2, 3 or 4 or more QC divisions. The number of analyzed roots for each genotype is indicated above each bar and result from to 3-5 technical replicates.

Additionally, the quantification of QC divisions was performed by counting the number of EdU-stained nuclei within an optical transversal section through the RAM as described in (Burkart *et al*., 2022). QC cells were identified by their relative position within the RAM, directly below the vascular initials and surrounded by CEIs in a circular arrangement (Fig. 2 K). In the WT, 57 % of the roots do not show any QC cell divisions, and 35 % show one QC cell division (Fig. 2 L, T, Fig. S1 A). In 6 % and 2 % of the analysed roots, two and three QC divisions could be observed, respectively, so that in total 43 % of the analysed roots showed EdU-stained QC cells. In *bravo*-2 and *plt3-1* single mutants, the number of roots showing at least one EdU-stained QC cell increased to 78 % and 73 %, respectively (Fig. 2 M, N, T, Fig. S1 B, C). This phenotype is even more severe in *wox5-1* mutants, where at least one EdU-stained QC cell could be observed in 86 % of the roots (Fig. 2 O, T, Fig. S1 D). Like the above-described additive effects of CSC differentiation in the double and triple mutants, the number of roots showing at least one QC cell division increases to 100 % and 98 % in the *bravo wox5* and *plt3 wox5* double mutants, respectively (Fig. 2 P-R, T, Fig. S1 E-G). Additionally, the double mutants show a strongly increased frequency of four divided QC cells in comparison to the respective single mutants: 7 % in the *bravo plt3* double mutant, 36 % in the *bravo wox5* double mutant and 30 % in the *plt3 wox5* double mutant in comparison to 3 %, 0 % and 11 % in the *bravo-2*, *plt3-1* and *wox5-1* single mutants, respectively. A further increase in EdU-stained QC cells can be observed in the *bravo plt3 wox5* triple mutant where 44 % of the roots display a completely divided QC (Fig. 2 S, T, Fig. S1 H). These observations indicate that BRAVO, PLT3 and WOX5 jointly control QC divisions, which may also involve other factors, e. g. SHORT-ROOT (SHR) and SCARECROW (SCR) (Cruz-Ramírez *et al*., 2013; Long *et al*., 2017; Clark *et al*., 2020).

Furthermore, we also examined if the QC exhibits extra periclinal cell divisions, which in *Col-0* WT occurs only in 4 % of the roots (Fig. S1 I, K). This phenotype manifests in 85 % of *bravo-2* mutants (Fig. S1 J, K). Additional periclinal cell divisions can also be observed in 43 % of *plt3-1* single mutants and in 62 % *wox5-1* single mutants (Fig. S1 K). In contrast to the number of EdU-stained QC cells, the frequency of periclinal cell divisions are relatively similar in the double or triple mutants, with 77 %, 84 %, 79 % and 85 % of the roots showing additional periclinal cell divisions of the QC cells in the *bravo plt3*, *bravo wox5*, *plt3 wox5* and *bravo plt3 wox5* mutants, respectively (Fig. S1 K). This effect has already been described for *wox5-1* and *bravo-2* single mutants in comparison to the *bravo wox5* double mutant in earlier studies (Betegón-Putze *et al*., 2021).

Taken together, our findings suggest a combinatory effect of BRAVO, PLT3, and WOX5 on QC division frequency and CSC fate decision.

### BRAVO, PLT3 and WOX5 can form a trimeric complex

In addition to the observed overlapping yet cell type specific protein levels and the genetic interplay of BRAVO, PLT3 and WOX5, recent reports also provide evidence for one-on-one PPIs of BRAVO and WOX5, as well as for PLT3 and WOX5 (Betegón-Putze *et al*., 2021; Burkart *et al*., 2022). These findings raised the question if also BRAVO and PLT3 could interact. To address this, we first performed fluorescence resonance energy transfer fluorescence lifetime imaging microscopy (FRET-FLIM) measurements in transiently expressing *Nicotiana benthamiana* (*N. benthamiana*) abaxial epidermal leaf cells using BRAVO-mV as donor molecule under control of a β-estradiol inducible promoter as described earlier (Burkart *et al*., 2022). Results of FRET-FLIM measurements are often displayed as the average amplitude-weighted lifetime which is a mixture of differentially decaying components and is calculated by summing each component’s lifetime weighted by its respective amplitude. In case of FRET, the fluorescence lifetime decreases and serves as a measure for PPI. This reduction of lifetime results either from a large number of molecules that undergo FRET indicating a high affinity of the two proteins of interest (POIs) or a highly efficient energy transfer which demonstrates high proximity of the POIs and/or favourable fluorophore dipole orientation (Fig. 3. A, B). The use of a novel analysis method allowed us to distinguish between these two scenarios, providing deeper insights into protein affinities, hereafter referred to as ‘binding’, between BRAVO, PLT3 and WOX5 (Maika *et al*., 2023).

**Figure 3:**
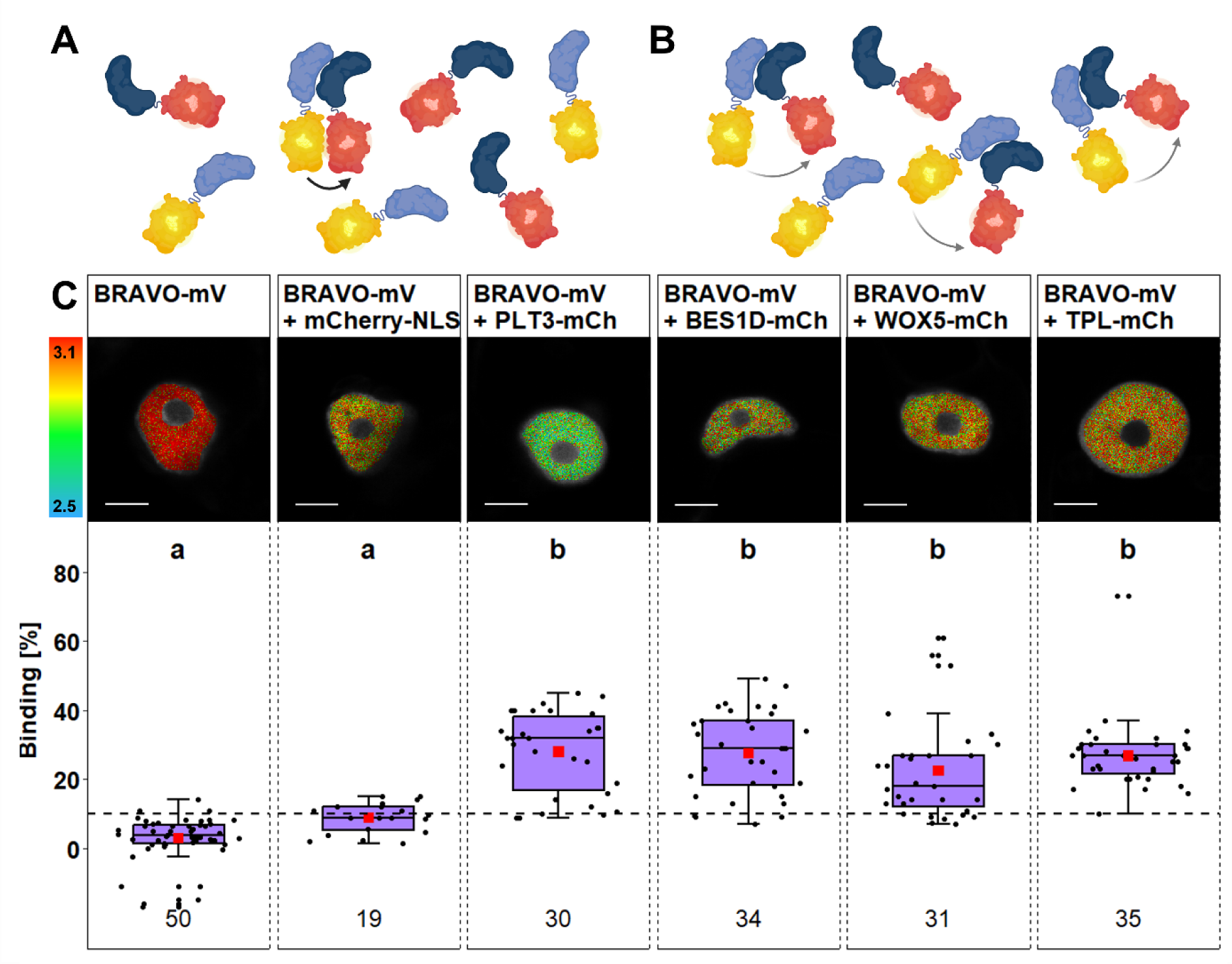
BRAVO interacts with PLT3, WOX5, BES1D and TPL. A) A reduction of fluorescence lifetime as a consequence of FRET can either be a result of a highly efficient energy transfer indicating close proximity or **B)** a high affinity of the two proteins. Figure created with BioRender.com and modified from (Maika *et al*., 2023). **C) Upper panel:** Representative images of fluorescence lifetime imaging microscopy (FLIM) measurements of nuclei in *N. benthamiana* epidermal leaf cells after pixel-wise mono- or biexponential fitting. The fluorescence lifetime of the donor BRAVO-mV in absence or presence of the indicated acceptor (of mCherry-NLS, PLT3-mCh, BES1D-mCh, WOX5-mCh or TPL-mCh) is color-coded: blue (2.5) refers to low fluorescence lifetime [in ns], red (3.1) indicates high fluorescence lifetime [in ns]. Scale bars represent 6 µm. **Lower panel:** Binding values [%] are represented as purple box plots of the same samples as in the upper panel. Statistical groups were assigned after a non-parametric Kruskal-Wallis ANOVA with *post-hoc* Dunn’s test (α = 0.05). Mean values are visualised as red squares. Black dotted line indicates the Binding cut-off of 10 %. Number of analysed nuclei is indicated below each sample and results from 3-5 technical replicates. Partially created with BioRender.com.

The reference sample BRAVO-mV (donor-only control) shows an average binding of 2.3 ± 7.4 % (Fig. 3 C) and the negative control composed of BRAVO-mV co-expressed with mCherry-NLS shows a binding of 8.8 ± 4.3 % (Fig. 3 C). A binding of below 10 % is interpreted as no interaction (Maika *et al*., 2023), cohering with the reference and negative control samples. Upon co-expression of BRAVO-mV with PLT3-mCh, the binding increases to 28.0 ± 11.7 % (Fig. 3 C). To compare this observation with already confirmed interactions of BRAVO with WOX5 (Betegón-Putze *et al*., 2021), as well as with BES1 or TPL (Vilarrasa-Blasi *et al*., 2014), we co-expressed BRAVO-mV with WOX5-mCh or TPL-mCh, which results in binding values of 22.4 ± 14.1 % and 26.7 ± 9.8 %, respectively (Fig. 3 C). Interaction of BRAVO with BES1 was tested by co-expression of BRAVO-mV with BES1D-mCh, which was shown to mimic the dephosphorylated and thereby active form of BES1 and yielded an average binding of 27.4 ± 11.9 %. This suggests similar affinities of BRAVO towards PLT3, BES1 and TPL, but a lower affinity towards WOX5 (Fig. 3 C).

These findings together with previously described interactions of WOX5 with PLT3, TPL or BES1, as well as BES1 and TPL, prompted us to investigate, whether these TFs can also form higher-order complexes (Vilarrasa-Blasi *et al*., 2014; Espinosa-Ruiz *et al*., 2017; Betegón-Putze *et al*., 2021; Burkart *et al*., 2022). To address this, we used a combination of bimolecular fluorescence complementation (BiFC) and FRET (Fig. 4 A, B) (Kwaaitaal *et al*., 2010; Maika *et al*., 2023). Here, the donor fluorophore is split into two fragments: the N-terminal part of mVenus (mV(N)) and the C-terminal part (mV(C)). The interaction of WOX5 and PLT3, which has been described earlier (Burkart *et al*., 2022), has been shown to have a high affinity (Supplemental table S13). This is why we have chosen to tag WOX5 and PLT3 with mV(N) and mV(C), respectively. In this scenario, the interaction of WOX5 and PLT3 leads to the reconstruction of mV and restores its fluorescence, enabling us to perform FRET-FLIM when co-expressing another acceptor-labelled protein. The ‘donor only’ reference sample WOX5-mV(N) PLT3-mV(C) yields an average binding of 1.6 ± 14.1 %, and the negative control WOX5-mV(N) PLT3-mV(C) with mCherry-NLS shows an average binding of 2.9 ± 5.0 % (Fig. 4 C). Upon co-expression of BES1D-mCh or TPL-mCh, the binding significantly increases to 18.7 ± 8.0 % and 23.3 ± 8.3 %, respectively (Fig. 4 C). Notably, in the presence of BRAVO-mCh, the average binding strongly increases to 36.3 ± 10.7 % (Fig. 4 C). Thus, the heterodimer of WOX5 and PLT3 shows higher affinity to BRAVO, which could suggest an increased probability and stability of the trimeric complex composed of WOX5, PLT3 and BRAVO compared to WOX5, PLT3 and BES1D, or TPL.

**Figure 4:**
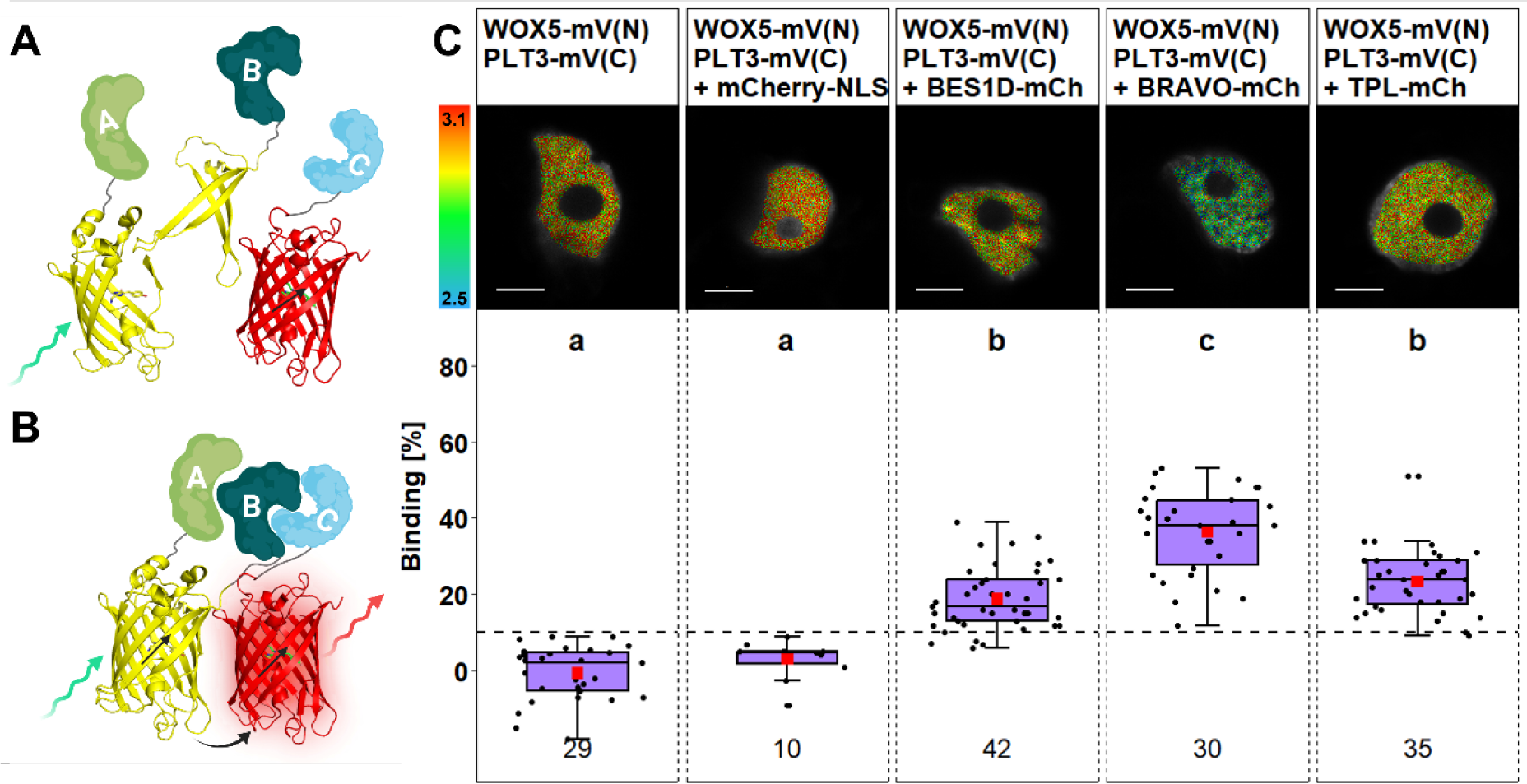
Trimeric complex formation of WOX5 and PLT3 with BRAVO, BES1D and TPL. A) The combination of BiFC-FRET allows the detection of higher-order complexes. Here, the two fragments of a split donor fluorophore are fused to two proteins of interest (POI), while a third POI is fused to the acceptor. **B)** In case of trimeric complex formation, the donor molecule is reconstructed and transfer energy to the acceptor molecule by FRET after excitation. Created with BioRender.com and modified from (Strotmann and Stahl, 2022). **C) Upper panel:** Representative images of fluorescence lifetime imaging microscopy (FLIM) measurements of nuclei *N. benthamiana* epidermal leaf cells after pixel-wise mono- or biexponential fitting. The fluorescence lifetime of the donor WOX5-mV(N)/PLT3-mV(C) in absence or presence of the indicated acceptor (mCherry-NLS, BES1D-mCh, BRAVO-mCh or TPL-mCh) is color-coded: blue (2.5) refers to low fluorescence lifetime [in ns], red (3.1) indicates high fluorescence lifetime. Scale bars represent 6 µm. **Lower panel:** Binding values [%] are represented as purple boxplots of the same samples as in the upper panel. Statistical groups were assigned after non-parametric Kruskal Wallis ANOVA with *post-hoc* Dunn’s test (α = 0.05). Mean values are visualised as red squares. Black dotted line indicates the Binding cut-off of 10 %. Number of analysed nuclei is indicated below each sample and results from 2-3 technical replicates. Partially created with BioRender.com.

To gain further insights into the potential of trimeric complex formation, we conducted additional FRET-FLIM measurements in *N. benthamiana* with rearranged fluorescent tags. Here, the donor fluorophore is shared between BRAVO and PLT3, namely BRAVO-mV(N) and PLT3-mV(C), which also showed high affinity (Fig. 3). The ‘donor only’ reference sample BRAVO-mV(N) PLT3-mV(C) exhibits an average binding of 2.2 ± 3.6 % (Fig. S2). The negative control composed of BRAVO-mV(N) PLT3-mV(C) and mCherry-NLS shows a similar average binding of 3.2 ± 3.1 % (Fig. S2). Surprisingly, co-expression of BES1D-mCh yields an average binding of only 10.2 ± 5.9 % (Fig. S2), indicating that a trimeric complex composed of BRAVO, PLT3 and BES1D is unlikely to form. Contrary, the co-expression of TPL-mCh or WOX5-mCh leads to a significantly increased average binding of 21.1 ± 8.3 % and 29.8 ± 10.6 %, respectively (Fig. S2). This again suggests that a trimeric complex formed by BRAVO, PLT3 and WOX5 is more stable and occurs with a higher probability. Taken together, these findings reveal the formation of several combinations of protein multimers with different probabilities of occurrence as judged by their binding capacities. Here, the complex composed of BRAVO, PLT3 and WOX5 seems to be the most frequent and stable.

### Modelling reveals cell type specific TF complex compositions

Our results reveal distinct, cell type specific patterns of protein abundance for BRAVO, PLT3 and WOX5 in the root SCN (Fig. 1) along with the formation of diverse heterodimers with varying binding affinities as well as higher-order complexes in *N. benthamiana* (Fig. 3, Fig. 4, Fig. S2). The protein complexes formed in the cells of the root SCN are ultimately a result of the cell type specific protein levels and the binding affinities between the proteins. This raises the question whether dimerization and complex formation in the context of the root apex also display cell type specificity, and how this is influenced by the protein levels in each cell of the SCN (Fig. 1). For example, BRAVO protein levels in the QC are notably lower compared to PLT3 or WOX5 (Fig. 1 H), yet its consequence on protein complex formation remains undetermined. While the FRET-FLIM approach could in theory be used to investigate the formation of dimer- and oligomerization in *Arabidopsis* root cells, previous efforts to assess the interaction of PLT3 and WOX5 under the control of their endogenous promoter in roots have been challenging due to limited protein abundance and, consequently, low photon counts (Burkart *et al*., 2022). This is a limitation difficult to overcome without altering the endogenous protein levels. Therefore, as an alternative to identify potential TF specificity and cell type specific complexes in the root SCN, we use a two-step mathematical modelling approach that combines the endogenous protein abundances (Fig.1) with the binding probabilities for one-on-one PPIs and trimeric protein complexes (Fig. 3, Fig. 4, Fig. S2).

First, we performed a parameter analysis to predict the relative association and dissociation rates to form the WOX5-PLT3, BRAVO-PLT3, BRAVO-WOX5 heterodimers, and the WOX5-PLT3-BRAVO trimeric complex. For the trimeric complex, we evaluate its formation via WOX5-PLT3 and BRAVO-PLT3 as donors (Fig. 4, Fig. S2). We start our simulations with equal levels of both donor and acceptor as initial condition, to mimic the conditions in the *N. benthamiana* experiments. Then, we simulate the protein complex formation using association and dissociation rates from a wide range of possible parameter values, until a steady state is reached. For each parameter combination tested, we evaluated if the proportion of protein in complex in steady state corresponds to the value from the respective relative binding affinity determined with our experiments. Repeating this parameter estimation for each of the protein complexes under study, allows us to identify several parameter combinations capable of producing protein complexes in line with FRET-FLIM experimental data (Fig. S3, Fig. S4). The predicted parameter combination for protein complexes with a high binding affinity (i.e. WOX5-PLT3) fall in the space where association rates are higher than the dissociation rates (Fig. S3), in contrast to lower binding affinity complexes (i.e. BRAVO-WOX5). These determined parameters allow us to describe our binding experimental data in a computational model.

Next, we simulated the protein complexes formed by BRAVO, WOX5 and PLT3 in each of the cells of the root SCN. For this, we use as initial condition the values from the relative fluorescence intensities we quantified for BRAVO, PLT3 and WOX5 in the SI, QC, CSC, and CC (Fig. 1 H), and the association/dissociation rates per complex from our parameter analysis. Therefore, the cell type specific profiles of protein complexes predicted by modelling are the emergent result of how much protein is available in each cell type and the binding affinities between specific protein pairs and complexes (Fig. 5). We summarized these results in a radar chart where the level of each protein complex is arranged in a different radial axis and displayed free protein levels that remain after complex formation separately as bar plots (Fig. 5 A, B). Furthermore, we combined these results in a heat map (Fig. 5 C). Additionally, we performed several controls that assume different combinations of experimental data, both binding affinities and cell type specific protein abundances, and varying ratios of association and dissociation (Fig. S5, Material and Methods). Interestingly, results comparable to our model were only observed in control 2, assuming higher association and dissociation rates, which indicates higher association also in our experimental data.

**Figure 5.**
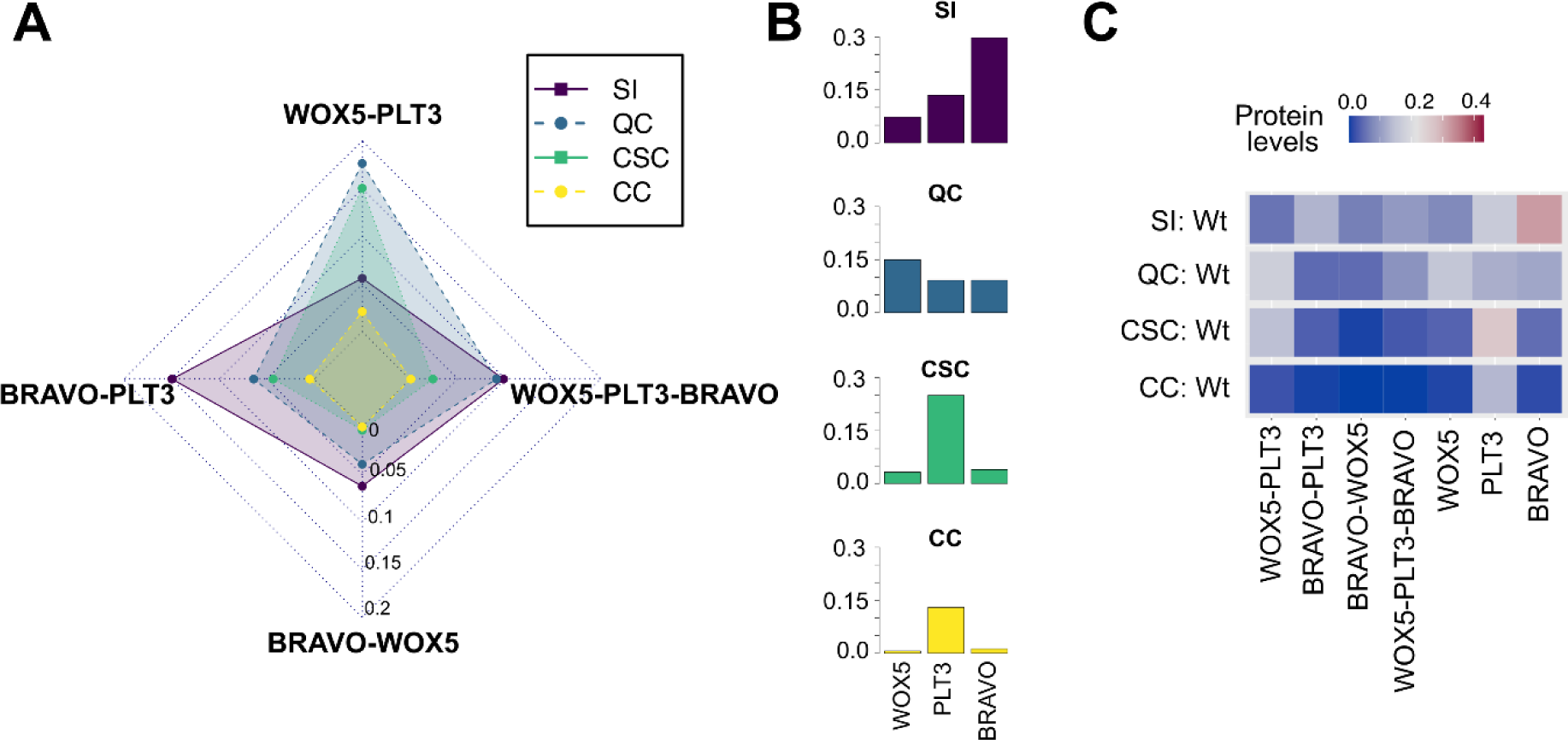
*In silico* prediction of protein complex signatures in the WT root SCN. A) Radar plot showing the levels of heterodimers and trimeric complex of WOX5, PLT3 and BRAVO formed in the SI (purple), QC (blue), CSC (green) and CC (yellow). The radial axis shows the protein levels (in arbitrary units). **B)** Free WOX5, PLT3 and BRAVO protein in each of the simulated root SCN cells. **C)** Heatmap showing the protein complexes and free protein in the cells of WT simulation. High concentrations are displayed in red, low concentration are displayed in blue. SI: stele initals; QC: quiescent center; CSC: columella stem cells; CC: columella cells.

Our simulation reveals that SIs are characterized by high levels of BRAVO-PLT3 protein complex (Fig. 5 A, C). The QC cells are predicted to be enriched in the WOX5-PLT3 complexes, followed closely by the CSC. Such enrichment could be related to the previously described function of the WOX5-PLT3 complex in QC divisions and CSC maintenance (Burkart *et al*., 2022). However, predictions of the trimeric complex WOX5-PLT3-BRAVO displays only intermediate levels in both the SI and the QC. Finally, the CCs are predicted to have negligible levels of all protein complexes studied, consistent with the very low BRAVO, PLT3, and WOX5 protein levels present in these cells according to our quantification (Fig. 1). Notably, these protein complex ‘signatures’ are strikingly different in each of the simulated cells and the resulting polygons are unique for each cell type (Fig. 5 A), which might be related to their specific function.

Curiously, the levels of free protein also show cell type specific patterns, that allow to further distinguish between SIs, QC and CSCs (Fig. 5 B, C). SIs are enriched in free BRAVO, while the QC shows high levels of free WOX5. Both, CSCs and CCs, exhibit high levels of PLT3. It is interesting to consider that these free proteins could participate in both, binding other proteins not considered here, and/or intercellular movement, assuming an increased mobility if the protein is not in complexes (Fig. 5 B, C). For instance, the levels of free WOX5 in the QC cells could constitute a pool of free protein available for intercellular mobility towards the neighbouring CSCs as previously described (Pi *et al*., 2015). In summary, these results support the hypothesis that complex formation, especially heterodimerization, occurs in a cell type specific context.

### Prion-like domains of PLT3 serve as conserved interaction hub

After we have found evidence for the formation of TF complexes with cell type dependent variations, we asked whether these complexes are important for root SCN maintenance. To address this, we aimed to destabilize the interaction of these TFs by mutating their specific interaction sites and observe if this altered protein can still rescue the phenotypical defects in the SCN. First, we explored the literature to identify potential interaction sites of BRAVO, PLT3 and WOX5. Previous studies have shown that prion-like domains (PrDs) in PLT3 mediate the interaction with WOX5 (Burkart *et al*., 2022). PrDs are intrinsically disordered regions (IDRs) and serve not only as mediators of multivalent interactions, but have also been demonstrated to be involved in chromatin opening (Levy *et al*., 2002) and phase separation (Jung *et al*., 2020). Given the presence of PrDs also in PLT1, PLT2 and PLT4, albeit in lower numbers (Burkart *et al*., 2022), we hypothesized that these regions function as conserved interaction sites. Thus, we performed FRET-FLIM measurements to investigate how the deletion of PLT3 PrDs, termed PLT3ΔPrD, affects its interaction with BRAVO. The ‘donor only’ reference control BRAVO-mV yields an average binding of 1.7 ± 4.6 %, which increases to 3.9 ± 2.4 % in the presence of mCherry-NLS serving as negative control (Fig. 6). Upon co-expression of BRAVO-mV with PLT3-mCh the binding significantly increases to 22.8 ± 10.5 % (Fig. 6). However, BRAVO-mV co-expressed with PLT3ΔPrD-mCh yields an average binding of only 11.7 ± 9.6 %, suggesting that the deletion of the PrDs significantly reduces the interaction of PLT3 with BRAVO.

**Figure 6:**
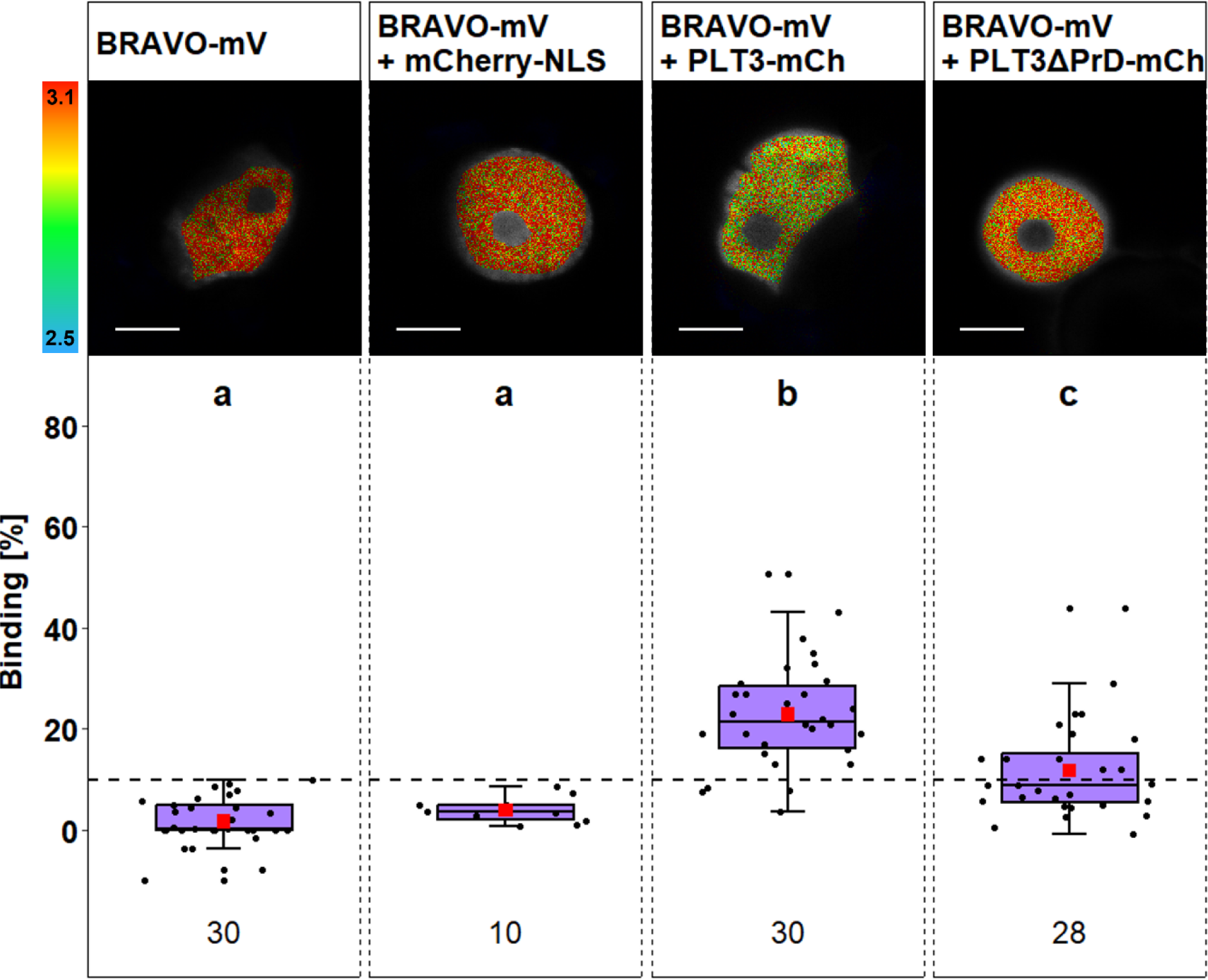
PrDs of PLT3 stabilize interaction with BRAVO. Upper panel: Representative images of fluorescence lifetime imaging microscopy (FLIM) measurements of nuclei in *N. benthamiana* epidermal leaf cells after pixel-wise mono- or biexponential fitting. The fluorescence lifetime of the donor BRAVO-mV in absence or presence of the indicated acceptor (mCherry-NLS, PLT3-mCh or PLT3dPrD-mCh) is color-coded: blue (2.5) refers to low fluorescence lifetime [in ns], red (3.1) indicates high fluorescence lifetime. Scale bars represent 6 µm. **Lower panel:** Binding values [%] are displayed as purple box plots of the same samples as the upper panel. Statistical groups were assigned after non-parametric Kruskal Wallis ANOVA with *post-hoc* Dunn’s test (α = 0.05). Mean values are visualised as red squares. Black dotted line indicates the Binding cut-off of 10 %. Number of analysed nuclei is indicated below each sample and results from 2-3 technical replicates.

To further support our hypothesis that the PrDs in PLTs act as conserved interaction site, we investigated whether PLT3 also interacts with BES1 and TPL, which were shown before for to interact with BRAVO and WOX5 (Vilarrasa-Blasi *et al*., 2014; Pi *et al*., 2015; Betegón-Putze *et al*., 2021) and if this interaction can also be altered by the deletion of PLT3 PrDs. To address this, we conducted FRET-FLIM in the presence of an acceptor-labelled PLT3 or PLT3ΔPrD. For the donor only reference control measurements BES1D-mV, an average binding of 0.0 ± 6.3 % could be observed which increases to 16.9 ± 8.1 % in the presence of PLT3-mCh indicating PPI (Fig. S6 A). However, co-expression of BES1D-mV with PLT3ΔPrD-mCh shows a reduced binding of 8.4 ± 4.8 % which is not significantly different from the negative control BES1D-mV with mCherry-NLS exhibiting an average binding of 4.9 ± 6.5 % (Fig. S6 A). The reference control TPL-mV exhibits an average binding of 0.6 ± 5.5 %, increasing to 6.4 ± 2.4 % when co-expressed with the negative control mCherry-NLS (Fig. S6 B). Upon co-expression of TPL-mV with PLT3-mCh, the average binding significantly increases to 13.5 ± 4.3 %, suggesting a moderate interaction of TPL with PLT3 (Fig. S6 B). Similar to BES1, the interaction of TPL and PLT3 is also abolished by the deletion of PrDs, demonstrated by a significantly decreased average binding of 8.99 ± 5.26 % for TPL-mV with PLT3ΔPrD-mCh (Fig. S6). In summary, these findings support the idea that the PrDs of PLT3 serve as a conserved interaction site for numerous TFs present in the root SCN.

### Redistribution of TF complexes alters regulation of QC divisions

Next, we aimed to analyse the functional relevance of the eliminated or reduced interaction of PLT3 with other TFs present in the *Arabidopsis* root by deleting its PrDs. To address this, we created two transgenic *Arabidopsis* lines, using either full-length PLT3 or PLT3ΔPrD C-terminally tagged with mTurquoise2 (mT2) in combination with the dexamethasone (DEX) inducible glucocorticoid receptor (GR) in the *plt3-1* mutant background. Using the *WOX5* promoter allowed us to specifically investigate how the loss of PLT3 PrD influences QC maintenance. These lines were named *pWOX5:GR-PLT3-mT2* (*pWOX5:iPLT3)* and *pWOX5:GR-PLT3ΔPrD-mT2* (*pWOX5:iPLT3ΔPrD)*. Finally, we performed a SCN staining and investigated if the QC exhibits additional periclinal cell divisions after inducing the plants by DEX treatment or in the presence of dimethyl sulfoxide (DMSO), which serves as a control (Fig. 7 A-I).

**Figure 7.**
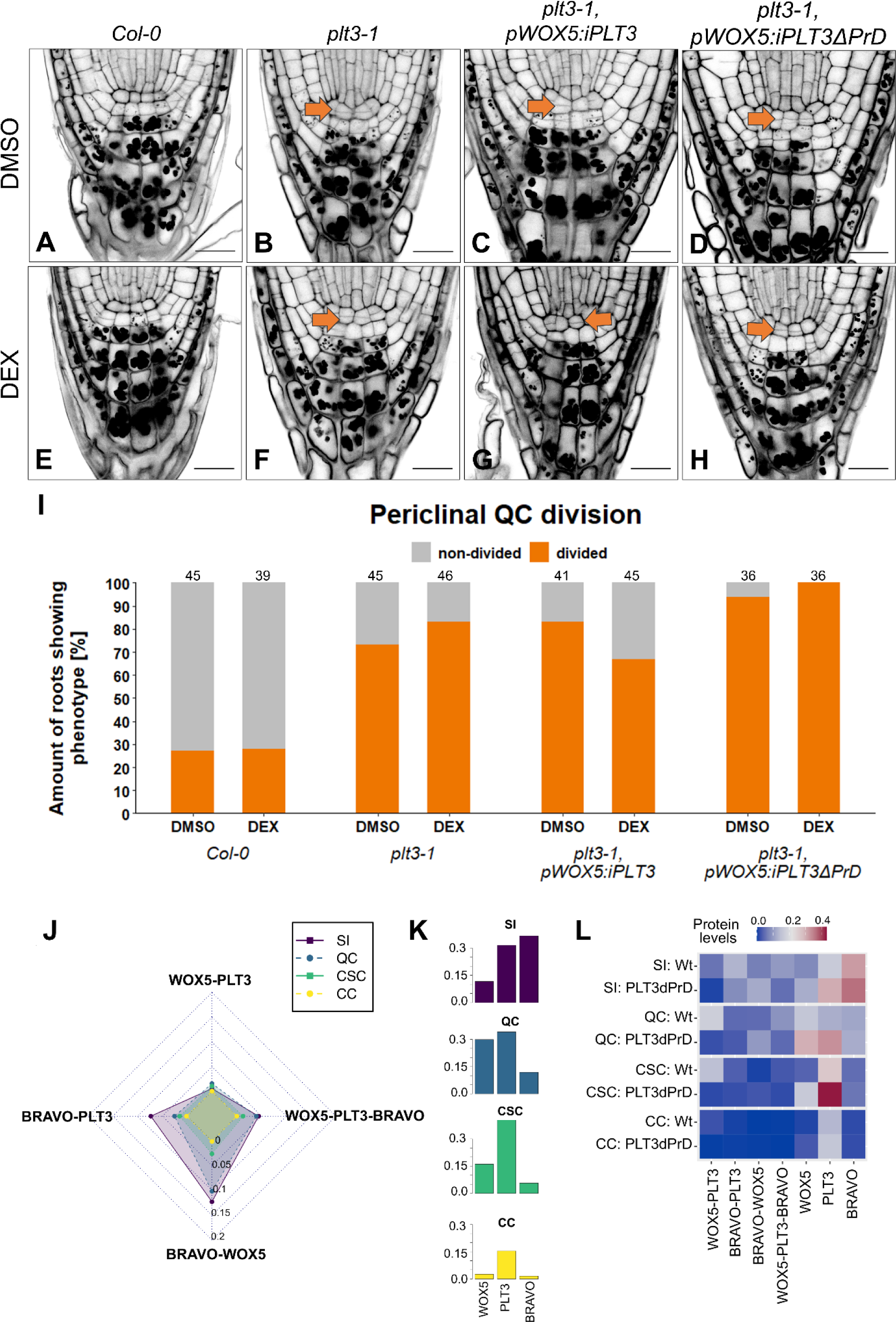
PLT3 PrDs inhibit periclinal QC divisions and *in silico* predicted protein complex signatures in the root SCN. Representative images of the *Arabidopsis* root meristem showing additional periclinal cell divisions in the QC in the absence **A-D)** or presence **E-H)** of DEX in the indicated genetic background. Divided QCs are highlighted with orange arrows. Scalebars represent 20 µm. **I)** Quantification of periclinal cell divisions when roots are treated with DMSO or DEX. Number of analysed roots is indicated above each bar and results from three replicates. **J)** Radar plot showing the levels of heterodimers and trimeric complex between WOX5, PLT3 and BRAVO formed in the stele initials (purple), QC (blue), CSC (green) and CC (yellow). **K)** Free WOX5, PLT3 and BRAVO protein in each of the simulated root SCN cells. **L)** Heatmap showing the protein complexes and free protein in the cells of WT and PLT3ΔPrD simulations, the profiles are visibly different with a marked increase in free PLT3 in the CSC. High concentrations are displayed in red, low concentration are displayed in blue. SI: stele initials; QC: quiescent center; CSC: columella stem cells; CC: columella cells.

Under control conditions, only 27 % of *Col-0* WT roots show additional periclinal cell divisions in the QC, which does not change significantly in the presence of DEX (Fig. 7 A, E, I). In agreement with previous observations (Burkart *et al*., 2022), *plt3-1* single mutant roots show additional periclinal cell divisions in the QC of 73 % under control conditions and 87 % after induction with DEX (Fig. 7 B, F, I). In *pWOX5:iPLT3* and *pWOX5:iPLT3ΔPrD* transgenic lines, 83 % and 94 % of the roots exhibit a periclinal cell division in the QC under control conditions, respectively, which is even higher than the *plt3-1* single mutant (Fig. 7 C, D, I). However, in the presence of DEX, only 67 % of the roots expressing *pWOX5:iPLT3* show this phenotype, indicating that full-length PLT3 in the QC partially restores the *plt3-1* periclinal cell division phenotype (Fig. 7 G, I). Contrary, the observed overproliferated phenotype that we see under control conditions in *pWOX5:iPLT3ΔPrD* mutant roots, is unaffected in the presence of DEX, indicating that the PrDs of PLT3 are necessary to inhibit additional periclinal QC divisions and thereby contribute to PLT3 function in root SCN maintenance (Fig. 7 H, I).

After observing the reduced affinity of PLT3ΔPrD for BRAVO and WOX5, and that it was unable to rescue SCN defects in *plt3-1* single mutants, we decided to use our computational model to predict immediate changes in the protein complex ‘signatures’ in the root SCN that may have contributed to this failed rescue. Thus, we simulated the protein complex formation in the SI, QC, CSC, and CC as described before but set the association rate of PLT3ΔPrD-WOX5 to zero (Burkart *et al*., 2022) and use the binding affinity we have determined experimentally for PLT3ΔPrD-BRAVO (Fig. 6 and S6). This leads to a dramatic shift in the protein complex ‘signatures’ of the root SCN cells (Fig. 7 J-L). The elimination of WOX5-PLT3 dimer formation causes a redistribution of PLT3 and WOX5 to the other protein complexes and an increase of free PLT3 and WOX5 protein levels in the SI and the QC cells, as well as higher levels of free PLT3 in the CSC (Fig. 7 J, K). While the BRAVO-PLT3 complex can still be formed, it is noticeably reduced in the SI, QC and CSC cells. Furthermore, the BRAVO-WOX5 complex levels increase in the SI and QC cells. Surprisingly, the profile of the trimeric complex shows only minor disruptions in the modelled cells. Therefore, even if the WOX5-PLT3 protein complex cannot be formed due to the removal of PLT3 PrDs, the trimeric complex can still be formed by the association of WOX5 with the BRAVO-PLT3 protein complex. Altogether, our PLT3ΔPrD simulation provides insights into the alterations on cell type specific protein levels that could be causative for defects observed experimentally in the root SCN.

## Discussion

In the past decades, our understanding of stem cell function and maintenance in the root of *Arabidopsis* has witnessed significant advances. Various aspects, including hormonal, developmental, as well as stress-related mechanisms have been discovered (Nolan *et al*., 2020; García-Gómez *et al*., 2021; Ubogoeva *et al*., 2021; Strotmann and Stahl, 2021). However, the underlying intricate network of molecular factors, still remains largely enigmatic. In this study, we aimed to unravel a new aspect of the regulatory network that controls root SCN maintenance, related to protein complex formation.

By utilizing a distinct SCN staining technique (Burkart *et al*., 2022), we assessed phenotypical defects in the architecture of the *Arabidopsis* root SCN of several single and multiple mutants (Fig. 2). We observed an increased CSC differentiation and an elevated periclinal QC division frequency in the SCN of *plt3-1* mutants (Burkart *et al*., 2022). The observed phenotypes agree with previous observations, and their relatively moderate phenotypic manifestation can be attributed to the substantial redundancy within the PLT TF family (Galinha *et al*., 2007; Burkart *et al*., 2022). Moreover, these observations are consistent with a uniform PLT3 protein abundance in SIs, QC and CSCs (Fig. 1). Compared to *plt3-1* single mutants, we could observe a stronger effect for QC division frequency in *bravo-2* single mutants but a similar mild phenotype for CSC differentiation. Again, these results are supported by the observed protein levels: Although BRAVO is most abundant in SIs, it can also be found in the QC, whereas it is notably reduced in CSCs. *wox5-1* single mutants show a severely defective root SCN, as demonstrated by the loss of CSCs and greatly increased periclinal QC divisions, as described before (Sarkar *et al*., 2007; Cruz-Ramírez *et al*., 2013; Pi *et al*., 2015; Betegón-Putze *et al*., 2021; Burkart *et al*., 2022). Similar to PLT3 and BRAVO, these phenotypes correlate with high WOX5 protein levels in the QC and less protein in the CSC where WOX5 was shown to move to (Pi *et al*., 2015; Berckmans *et al*., 2020).

In the *bravo plt3*, *bravo wox5*, and *plt3 wox5* double mutants, we observed an increase in both QC division frequency and CSC differentiation, that were consistently higher than the respective single mutants. For PLT3 and WOX5 such additive effects have been described before and were hypothesized to show that they act in parallel pathways to maintain the integrity of the root SCN (Burkart *et al*., 2022). However, previous findings suggest that BRAVO and WOX5 act in the same pathway to control CSC fate and QC divisions based on quantifications of additional periclinal cell divisions (Betegón-Putze *et al*., 2021). We could observe similar effects when analysing periclinal cell divisions in the QC but using a novel SCN staining technique, we observed additive effects for QC division alterations in the *bravo wox5* double mutant compared to the respective single mutants (Fig. 2, Fig. S1). Our findings suggest the presence of an additional pathway that involves BRAVO and PLT3. Moreover, this indicates that these TFs could act in three independent constellations to regulate SCN maintenance. However, in the *bravo plt3 wox5* triple mutant, an additional additive effect could only be observed for QC divisions but not for CSC differentiation. A potential interpretation of these results is that none of these TFs is involved in an additional pathway to control CSC differentiation. However, they may partially contribute to other pathways that inhibit QC divisions. Additional functions in other independent pathways have already been described for WOX5 in the SHR-SCR regulatory network (Cruz-Ramírez *et al*., 2013; Zhai *et al*., 2020; Clark *et al*., 2020). Additionally, TEOSINTE-BRANCHED/CYCLOIDEA/PCNA 20 (TCP20) was found to mediate the interaction of PLT3 and SCR, to specify the QC and establish the root SCN (Shimotohno *et al*., 2018). If and to what extent these molecular factors genetically interact with other TFs in the SCN, will be an interesting perspective for future investigations.

In addition to the identified genetic interplay of BRAVO, PLT3 and WOX5 regarding root SCN maintenance, we were able to evaluate their physical interaction (Fig. 3, Supplementary Table S13). While interactions of PLT3 and WOX5 as well as BRAVO and WOX5, have been described before (Betegón-Putze *et al*., 2021; Burkart *et al*., 2022), evidence for an interaction of PLT3 and BRAVO was still missing. Our results reveal for the first time PPI between BRAVO and PLT3 as well as between PLT3 and BES1 and TPL (Fig. 3, Fig. S6). Together with previously described, independent one-on-one interactions, these findings support the hypothesis of three parallel pathways that control CSC differentiation and QC divisions in parallel. Furthermore, the observed variations of stability and probability of occurrence as indicated by a special analysis tool (Orthaus *et al*., 2009; Maika *et al*., 2023), could indicate a specific mechanism that facilitates the interaction of two POIs in a highly dynamic microenvironment, where the number of proteins is generally high, such as in the QC. (Fig. 1).

Next, the combination of BiFC and FRET allowed us to investigate the formation of higher-order complexes (Fig. 4, Fig. S2). Like in the one-on-one interaction studies, we found differences in protein affinities of the complexes under investigation. Here, the trimeric complex formed by WOX5-PLT3-BRAVO appeared to be the most abundant and stable. The heterodimerization of transcriptional regulators increases binding specificity and affinity and allows the combination of different internal as well as external signal inputs into gene regulation (Strader *et al*., 2022). This idea is reinforced when considering that both the auxin-regulated WOX5 and BR-dependent BRAVO have been demonstrated to control the same cell cycle-related genes (*CYCD1;1, CYCD3;3*) (Forzani *et al*., 2014; Vilarrasa-Blasi *et al*., 2014). So far, cell cycle-related downstream targets of PLT3 remain unknown. Further investigations are necessary to uncover potentially common downstream targets of BRAVO, PLT3 and WOX5.

To elaborate on differences in protein abundance and complex formation in cells of the root SCN, we used a computational modelling approach. This strategy allowed us to describe cell type specific protein complex profiles in WT roots (Fig. 5). Here, the combination of high levels of the BRAVO-PLT3 heterodimer and high levels of free BRAVO appears to be characteristic for stele initials. Interestingly, BRAVO protein abundance not only decreased when moving distally from the SIs, but also in proximal direction (Fig. 1). However, alterations of SCN defects in *bravo-2* single mutants had only been evaluated for CSC differentiation and QC division. New phenotypical analyses are necessary to determine whether SIs and their descendants are also affected upon loss of BRAVO function.

Our simulations of protein ‘signatures’ revealed that both, QC as well as CSC, are enriched in the WOX5-PLT3 heterodimer, which aligns with their previously described impact on QC divisions and CSC differentiation (Burkart *et al*., 2022). However, the protein ‘signatures’ of QC and CSC could be distinguished when free protein levels were considered. In the QC, our model predicted high protein levels of free WOX5, while CSCs were predicted to possess higher levels of PLT3. Several studies highlighted the elevated abundance of WOX5 in the QC, which could be either linked to interactions with other proteins not analysed here or its non-cell autonomous function in the adjacent initials, although its necessity as mobile stemness factor is still under debate (Pi *et al*., 2015; Berckmans *et al*., 2020). The predicted high levels of PLT3 protein in CSCs might be linked to nuclear body (NB) formation of PLT3, which was linked to its PrDs and is concentration dependent and may involve PLT3 homomerization. This mechanism could facilitate the recruitment of the WOX5-PLT3 heterodimer into these pre-formed NBs, as demonstrated previously (Burkart *et al*., 2022).

In CCs, the absence of BRAVO and WOX5 hinders complex formation, resulting in high levels of free PLT3. However, compared to CSC PLT3 levels are notably lower accompanied with loss of NBs formation. This implies that a specific protein concentration is required to initially trigger NB formation highlighting the difference between differentiated CCs and the stem cell fate determination process in CSCs. Based on our results, we created a final model that summarizes the described protein ‘signatures’ (Fig. 8). Here, SIs are characterized by high levels of free BRAVO protein and the heterodimer BRAVO-PLT3. QC cells and CSCs possess elevated levels of the WOX5-PLT3 heterodimer, which is accompanied by high levels of free WOX5 in the QC and high levels of free PLT3 in CSC. In CCs, complex formation is hindered by negligible levels of BRAVO and WOX5, resulting in elevated levels of free PLT3. All together our findings imply the formation of dimers that together with differences of free protein levels convey cell type specificity in the root. In the future, it should be addressed how the predicted protein complex ‘signatures’ drive changes in gene expression, including *BRAVO*, *PLT3*, and *WOX5*, but also other target genes, and how this relates to QC division and CSC number alterations in single and multiple mutants. As a next step, the model could also consider the complex gene regulatory networks in the root SCN (Cruz-Ramírez *et al*., 2012; García-Gómez *et al*., 2017; Pardal and Heidstra, 2021), the role of cell-cell mobility of free protein (Mähönen *et al*., 2014; Pi *et al*., 2015; García-Gómez *et al*., 2020; Betegón-Putze *et al*., 2021), the presence of membrane-less compartments to account for the localization of WOX5-PLT3 in nuclear bodies in the CSC (Burkart *et al*., 2022) and other key regulatory processes involved. The integration of experimental and computational approaches holds promise to uncover these complex mechanisms underlying root SCN maintenance.

**Figure 8:**
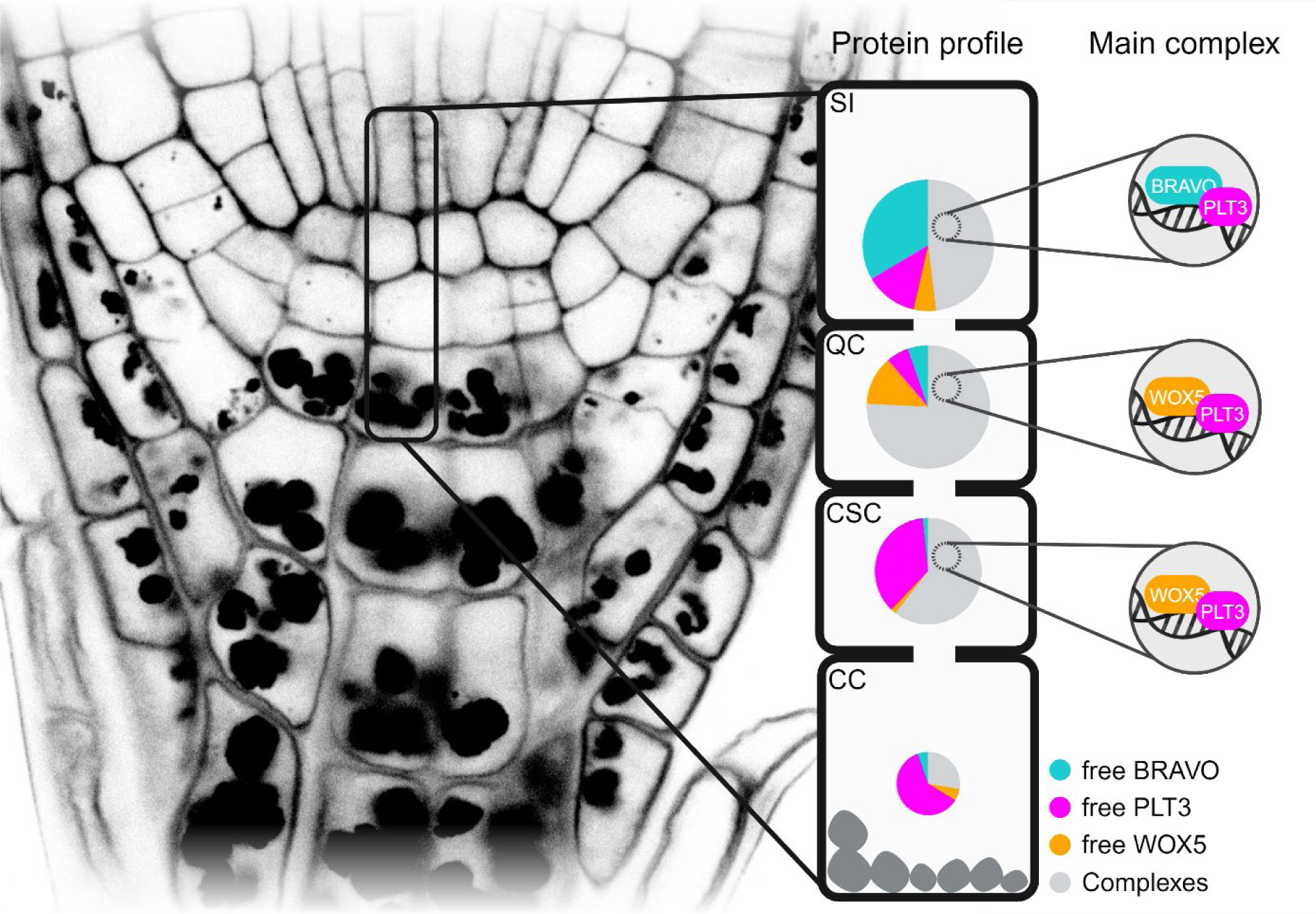
Model of protein signatures and complexes in the root SCN. The nuclei of different cell types (SI, QC, CSC, CC) show distinct protein profile of free BRAVO (turquoise), PLT3 (magenta), and WOX5 (orange) protein levels and main complexes (gray and insets). The size of the pie chart reflects the overall protein concentration in the nuclei of the specific cell type from high concentration (big) to low concentration (small). Created with BioRender.com.

To investigate the impact of heterodimer- and oligomerization on root SCN maintenance, we aimed to identify potential interaction sites in the BRAVO, PLT3 or WOX5 amino acid sequence. Previous studies revealed that PrDs found in PLT3 act as mediator of its interaction with WOX5 (Burkart *et al*., 2022). PrDs are also present in PLT1,2 and 4 which is also accompanied by NB formation. However, PLT3 harbours the highest number of PrDs, which correlates with stronger NB formation compared to PLT1, 2 and 4. Here, we demonstrated that loss of PrDs also negatively influences PLT3 interaction with BRAVO, BES1 and TPL (Fig. 6, Fig. S6). These findings suggest that PrDs act as a multivalent interaction hub, which could also indicate a conserved function among other PLTs.

In a rescue experiment, we could demonstrate that the PrDs of PLT3 affect its ability to inhibit periclinal QC divisions by demonstrating that PLT3ΔPrD, expressed in the QC, is unable to rescue the *plt3-1* periclinal QC division phenotype (Fig. 7). This indicates that correct dimer- and oligomerization is necessary for proper QC maintenance. We integrated our findings of diminished interactions of PLT3ΔPrD with BRAVO and WOX5 to our model and found a severe shift of protein complex ‘signatures’, especially for the WOX5-PLT3 dimer in the QC and CSCs. This further strengthens our hypothesis that the protein complexes form instructive protein signatures important for cell fate decisions in the *Arabidopsis* root SCN. Interestingly, full-length PLT3 under control of the *WOX5* promoter only partially rescues the *plt3-1* periclinal QC division phenotype. This emphasizes that functional PLT3 is also necessary to locally maintain CSC fate and repress differentiation as the QC divides to replenish lost CSCs (Cruz-Ramírez *et al*., 2013). Furthermore, this could indicate a specific function for PLT3 in the CSC fate, as the presence of other PLTs was not able to fully compensate for the loss of PLT3. Previous findings in yeast suggest that differences in IDRs mediate specificity of transcription factors that share the same DNA-binding motif (Brodsky *et al*., 2020). This is often observed among TFs that belong to the same family. If a similar mechanism also exists in plants, this could suggest that PLT3 function in CSC fate is specifically linked to its PrDs and that, due to their differentially structured PrDs, the other PLTs cannot compensate for this specific function. Additionally, this could indicate that mobile PLT3 which might move from the QC to CSC is not enough to maintain CSC stem cell character.

IDRs or PrDs also play a role in a recently described alternative mechanism of how TF find and locate to their specific DNA target (Staller, 2022). Indeed, the majority of TF found in eukaryotes is mainly composed of IDRs and only a small fraction of the protein sequence is well-characterized (Ward *et al*., 2004; Wang *et al*., 2016). According to this theory, IDRs of TFs scan the genome for matching protein clouds which mediate binding of the DNA-binding domain to its specific genomic target site (Staller, 2022). TFs possess two main functions: bind other TFs and bind to their specific DNA target to alter gene expression (Strader *et al*., 2022). Some TFs possess an additional important role; pioneer transcription factors, like LEAFY (Lai *et al*., 2021; Jin *et al*., 2021), bind to nucleosome bound DNA, open the target locus, e.g. by displacing H1 linker histones and/or recruiting chromatin remodellers, and make it accessible for other TFs. In plants, the concept of pioneer transcription factors is a newly emerging research field, but studies in animals suggest, that ‘master regulators’ appear to be promising candidates for pioneer transcription factors (as reviewed in Yamaguchi, 2021). The identification of IDRs and/or PrDs, that possess the ability to facilitate multivalent interaction and have been shown to act in chromatin opening (Levy *et al*., 2002), together with high redundancy within the PLT TF family, their role as master regulators of root formation and the stable protein abundance in the SCN, especially in cells that possess stem cell character, could indicate that also PLTs act as pioneer transcription factors in the root SCN. Interestingly, DNA affinity purification-sequencing (DAP-seq) results found PLT3, as well as PLT7, to be highly enriched in mCG-methylated DNA, providing yet another hint for this theory (O’Malley *et al*., 2016). Nevertheless, more evidence is necessary to further support the potential function of PLTs as pioneer transcription factor in root SCN maintenance.

Overall, our results suggest that BRAVO, PLT3 and WOX5 form cell type specific profiles of protein complexes and that proper complex formation contributes to optimal stem cell maintenance. Furthermore, we propose that these unique protein complex signatures serve as a read-out for cell specificity and could explain the different roles played by BRAVO, PLT3 and WOX5 in the regulation of stem cell homeostasis in the root.

## Material and Methods

### Plant work

All *Arabidopsis thaliana* lines used in this study were in *Col-0* background and can be found in Appendix Table S5. The *wox5-1* and *plt3-1* single mutants (Galinha *et al*., 2007) as well as the *bravo-2* single mutant (Vilarrasa-Blasi *et al*., 2014) and *bravo-2 wox5-1* double mutant (Betegón-Putze *et al*., 2021) were described before. The *bravo-2 plt3-1* double and *bravo-2 plt3-1 wox5-1* triple mutants were created by crossings. Homozygous F3 plants were verified by PCR using appropriate primers (Appendix Table S2). Transgenic lines were created by the floral dip method (Zhang *et al*., 2006). The *pPLT3:PLT3-mV* and *pWOX5:WOX5-mV* translational reporters in *Col-0* WT background were described earlier (Burkart *et al*., 2022). For *pBRAVO:BRAVO-mVenus*, *pWOX5:GR-PLT3-mTurquoise2,* and *pWOX5:GR-PLT3ΔPrD-mTurquoise2* transgenic plants, lines were selected, that possess a single T-DNA insertion, which was tested by observing the segregation on selection marker containing plates. Plants for crossing, genotyping, transformation, floral dip and amplification were grown under long-day conditions (8 h dark, 16 h light) at 21 °C and 60 % humidity. For microscopy, seeds were sterilized with chlorine gas (50 ml 13 % sodium hypochlorite (v/v), 1 ml hydrochloric acid) in a desiccator, mounted in 0.15 % (w/v) agarose and stratified in the dark at 4 °C for minimum two days before sowing on GM agar plates without sucrose (1/2 MS including Gamborg B5 vitamins, 1.2 % plant agar (w/v) and 0.05 % MES hydrate (w/v)). Seedlings for imaging were grown for five to six days under continuous light at 80 µmol m^-2^ s^-1^, 21 °C and 60 % humidity.

### Cloning

Plasmids for the transgenic lines *pBRAVO:BRAVO-mVenus*, *pWOX5:GR-PLT3-mTurquoise2* and *pWOX5:GR-PLT3ΔPrD-mTurquoise2* as well as for transient expression in *N. benthamiana* were generated using the GreenGate cloning method in the pGGZ001 destination vector (Lampropoulos *et al*., 2013). The region of the *WOX5* promoter, the CDS of WOX5, PLT3 and PLT3ΔPrD CDS as well as WOX5, PLT3 and PLT3ΔPrD constructs for transient expression in *N. benthamiana* were described before (Burkart *et al*., 2022). The region upstream of the transcriptional start of BRAVO (2,925 bp) (Lee *et al*., 2006) was assigned as promoter and amplified by PCR with appropriate primers containing flanking *Bsa*I restriction sites and matching overlaps for GreenGate cloning. The internal *Bsa*I recognition site in the *BRAVO* promoter region was not removed, but incubation times for restriction digestion and GreenGate reaction were adapted accordingly. After PCR, the promoter sequence was cloned into the GreenGate entry vector pGGA000 using *Bsa*I restriction and ligation. The CDS of BRAVO and TPL were amplified from cDNA derived from extracted RNA by PCR using primers carrying the *Bsa*I recognition site and matching GreenGate overhangs. Next, they were cloned into the GreenGate entry vector pGGC000 via restriction digest and ligation. All entry vectors were confirmed by sequencing. The GreenGate entry vector carrying the β-estradiol inducible promoter cassette was provided by (Denninger *et al*., 2019). For bimolecular fluorescence complementation, the GreenGate M and N intermediate vectors, each of which carried one expression cassette, were used. The correct assembly of the modules was confirmed by sequencing. All module combinations, constructs as well as primers used for cloning are listed in Appendix Tables S4, S3, and S1, respectively.

### SCN staining

SCN staining was performed according to (Burkart *et al*., 2022). For CSC layer quantification, optical longitudinal sections of the *Arabidopsis* root were acquired. The cell layer below the QC was scored as differentiated if three or more cells in this layer accumulated starch granules. QC cell divisions were quantified using an optical cross-section of the RAM on a scale of zero to four or more cells. If the QC was duplicated and showed two layers, as often seen for *bravo-2* mutants, only QC divisions in the upper layer were counted.

The CSC layer and QC cell division phenotypes were visualized separately in bar plots using Microsoft Excel (Microsoft Office 365, Microsoft Corporation). To assess potential correlations between CSC layers and QC divisions, data were combined into 2D-plots showing QC division on the x-axis and CSC layer on the y-axis using Origin 2021b (OriginLab Corporation).

### Transient expression in *Nicotiana benthamiana*

For transient expression in *N. benthamiana*, the *Agrobacterium* strain GV3101::pMP50 was used that in addition to the plasmid harbouring the desired construct, carried the helper plasmid pSOUP needed for GreenGate vectors. *Agrobacteria* were grown overnight in 5 ml dYT medium at 28 °C with shaking. After centrifugation for 10 min at 4,000 rpm and 4 °C, the pellet was resuspended in infiltration medium (5 % sucrose (w/v), 0.01 % MgSO_4_ (w/v), 0.01 % glucose (w/v) and 450 µM acetosyringone) to an optical density OD_600_ of 0.6 and mixed with an *Agrobacterium* strain carrying the p19 silencing repressor and eventually with a second *Agrobacterium* strain carrying a different construct for co-expression. Subsequently, the cultures were incubated for 1 h at 4 °C. To trigger stomatal opening and thereby allow easy infiltration, *N. benthamiana* plants were sprayed with water and kept under high humidity prior to infiltration. The abaxial side of the leaf was infiltrated using a syringe without a needle. Expression was induced 2-4 days after infiltration by spraying a 20 µM β-estradiol solution containing 0.1 % Tween®-20 (v/v) to the abaxial side of the leaf. Depending on the expression level, FLIM measurements were performed 2-16 h after induction.

### Microscopy

Imaging of *Arabidopsis thaliana* roots was performed using an inverted ZEISS LSM780 or LSM880. For cell wall staining, Arabidopsis seedlings were mounted in an aqueous solution of propidium iodide (PI) (10 µM). Fluorophores and fluorescent dyes were excited and detected as follows: PI was excited with 561 nm and detected at 590-670 nm; Alexa Fluor® 488 was excited at 488 and detected at 500-580 nm; mVenus was excited at 514 nm and detected at 520-570 nm and mCherry was excited at 561 nm and detected at 580-680 nm. When mVenus was co-expressed with mCherry, it was excited at 488 nm and detected at 505-555 nm.

### Intensity measurements of protein levels in *A. thaliana*

For analysis of expression levels of different reporters in 6 DAG *Arabidopsis* roots of different genotypes, an inverted LSM880 microscope with constant settings for all reporters was used. The mean fluorescence levels were measured in ImageJ using an oval region of interest (ROI) of the size of one nucleus. One to three nuclei were measured per cell type and root of which the mean was calculated. Data were normalized to mean value of the combination cell type and reporter that yielded the highest intensity. Data result from three technical replicates.

### Induction of GR inducible Arabidopsis lines

For the *plt3-1* rescue experiments, seeds were sown on GM agar plates without sucrose (1/2 MS including Gamborg B5 vitamins, 1.2 % plant agar (w/v) and 0.05 % MES hydrate (w/v)) containing either 0.1 % DMSO (v/v) for control condition or 20 µM DEX (diluted in DMSO) for GR induction. After 5 days, seedlings were transferred to GM agar plates without sucrose containing 7 µg/ml 5-ethynyl-2’- deoxyuridine (EdU) and either 0.1 % DMSO (v/v) or 20 µM DEX (diluted in DMSO) and grown for 24 h. SCN staining, imaging and scoring of QC divisions and CSC layers were performed as described above.

### FRET-FLIM measurements

FRET-FLIM measurements were performed in transiently expressing epidermal leaf cells of 3 to 4 weeks old *N. benthamiana* using an inverted ZEISS LSM 780 equipped with additional time-correlated single-photon counting devices (Hydra Harp 400, PicoQuant GmbH) and a pulsed laser diode. mVenus was chosen as donor and excited at 485 nm with 1 µW laser power at the objective (40 x C-Apochromat/1.2 Corr W27, ZEISS) and a frequency of 32 MHz and detected using two τ-SPAD single photon counting detectors in perpendicular and parallel orientation. Photons were collected over 40 frames at 256x256 pixels per frame, a pixel dwell time of 12.6 µs and a digital zoom of 8. Prior to image acquisition, a calibration routine was performed. To test system functionality, fluorescence correlation spectroscopy (FCS) measurements of deionized water and Rhodamine110 were acquired. Additionally, monitoring the decay of erythrosine B in saturated potassium iodide served as instrument response function (IRF) to correct the fitting for system specific time shift between laser pulse and data acquisition. First, fluorescence decays of the donor-only control were analysed using the ‘Grouped FLIM’ analysis tool to determine the average fluorescence lifetime using a mono- or biexponential fitting model (SymPhoTime, PicoQuant GmbH). Next, to extract information about protein affinities and proximities, the ‘Grouped LT FRET Image’ tool was utilized for a monoexponentially decaying donor and the ‘One Pattern Analysis (OPA)’ tool was used for samples with a biexponentially decaying donor (SymPhoTime, PicoQuant GmbH). These tools allow separate analyses of the amplitude and fluorescence lifetime of the FRET fraction of each sample. Consequently, the amplitude of the FRET component serves as a measure for the number of molecules undergoing FRET, termed binding or protein affinity, whereas the difference of the fluorescence lifetime of the FRET component compared to the lifetime of the donor-only fraction is used to calculate the FRET efficiency which serves as a measure for protein proximity and orientation (Maika *et al*., 2023). For samples where molecules do not undergo FRET e.g., the donor-only and negative control, binding values mostly varied between - 10 and 10 % and corresponding FRET efficiencies mostly accumulated at 10 or 80 %, which was defined during the fitting process (Maika *et al*., 2023).

### Statistical tests

Data were tested for normal distribution by Shapiro test (α = 0.05) followed by a Levene’s test for equality of variances (α = 0.05). Since some data did not show normal distribution or equality of variances or both, all data sets were tested with a non-parametric Kruskal-Wallis ANOVA with *post-hoc* Dunn’s test (α = 0.05). Statistical testing was performed using R.

#### Protein complex modelling

To estimate the relative association and dissociation rates for each of the dimeric and trimeric complexes studied here, we used the following ordinary differential equations:

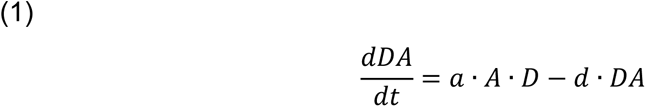

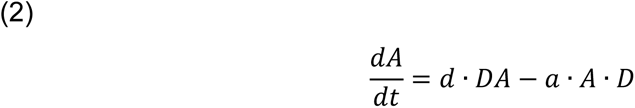

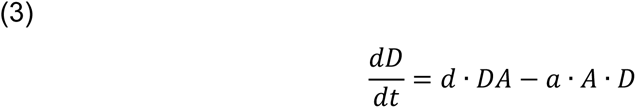

where *DA* is the protein complex formed by donor protein *D* and acceptor protein *A*. Using these equations, we simulated that the amount of protein complex, *DA*, is determined by the product of the association rate (*a*), the concentrations of donor, *D*, and acceptor, *A*, proteins, and how much it dissociates given a certain dissociation rate (*d*). To explain the relative binding affinity values determined experimentally for each dimeric and trimeric protein complex, we assessed association and dissociation rates involved in the protein complex formation from a wide range (0 – 0.5 arbitrary units, step 0.001), and simulated the protein complex *AB* formation until a steady state was reached. We deemed a particular combination of association and dissociation rates successful if they produce a value of *AB* at steady state in line with the relative binding affinity rates. In this way, we were able to predict relative binding rates for the dimeric and trimeric protein complexes studied here.

Next, we simulated the protein complex formation in the cells of the root SCN using the following ordinary differential equations to describe the formation of each dimeric and trimeric complex:

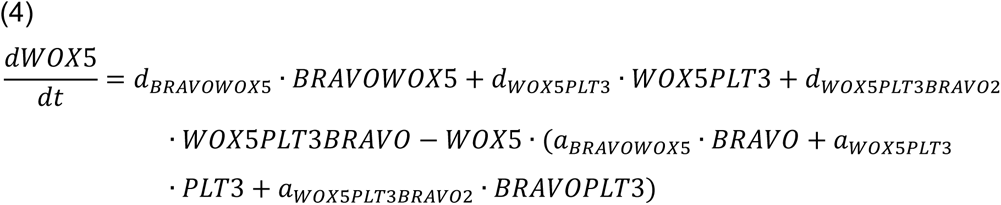

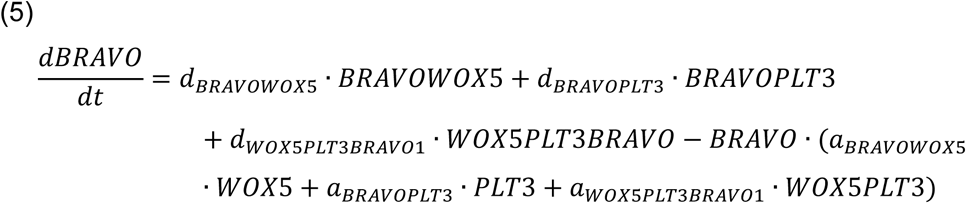

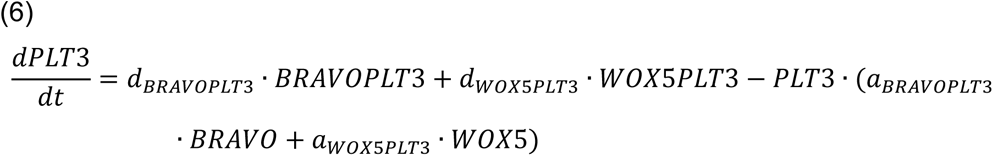

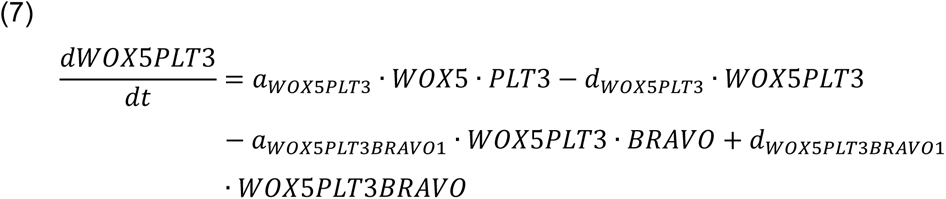

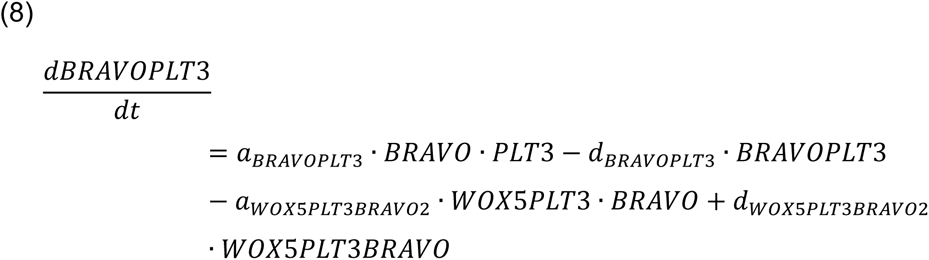

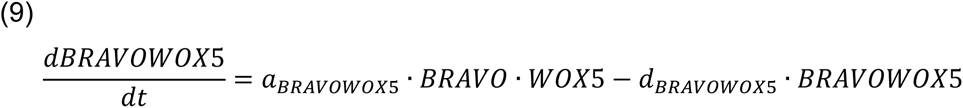

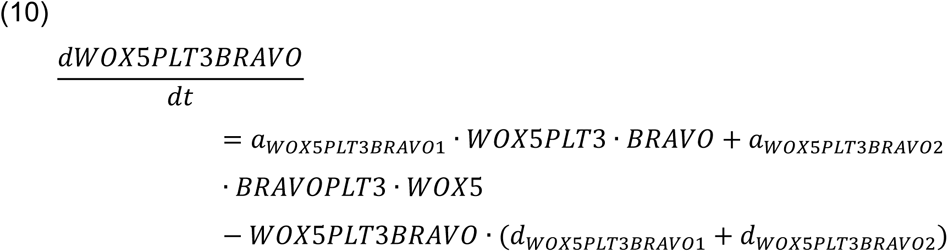

Notice the trimeric complex can be formed either by the binding of BRAVO to WOX5-PLT3, or WOX5 to BRAVO-PLT3. Then, we modelled the protein complexes formed in the cells of the root SCN using equations 4-10 and the relative protein levels of WOX5, BRAVO and PLT3 determined for SI, QC, CSC, and CC cells as initial condition. As several sets of binding rates were predicted per complex, for these simulations we used one selected at random. Notably, the specific parameters used for the results we present here do not change the protein complex signatures predicted by the model (Fig. S4).

To evaluate the effect in our model of both, the cell type specific protein levels as well as differential binding affinities are necessary for our model, we performed different control simulations. On the one hand, we tested the effect of equal association/dissociation rates (*a* = *d* = 0.1), higher association than dissociation rate (*a* = 0.1, *d* = 0.05), and lower association than dissociation (*a* = 0.05, *d* = 0.1) for all protein complexes using our experimental protein level quantification in the SI, QC, CSC and CC displayed as Control 1-3, respectively (Fig. S5). On the other hand, we consider an alternative scenario where all proteins have the same abundance levels, while the association/dissociation rates are based on our binding data (Control 4, Fig. S5). Finally, we consider the scenarios where the control conditions meet pairwise: Control 5 is a combination of equal association/dissociation rate together with the assumption of equal protein abundances among cell types and proteins. In Control 6, the equality of protein levels is combined with higher association than dissociation rates. Finally, Control 7 combines lower association that dissociation rates with equal protein abundances (Fig. S5). Notably only control 2, which uses experimentally determined protein abundances together with a higher association than dissociation rate, produced results comparable to our model. Thus, leading to the conclusion that also in our experimental data, association rates must be higher than dissociations rates. Moreover, this indicates a key role of the protein levels in each cell in the resulting protein complex and free protein signatures. In all other cases, we could observe strikingly different protein complex ‘signatures’ to those we described with the model that uses our experimental data, indicating that our findings result from the combination of experimentally determined specific protein levels and binding affinities.

The code for the computational model generated in this study was implemented in R, and will be available at the Garcia Group webpage in the server of the Theoretical Biology and Bioinformatics Group (https://bioinformatics.bio.uu.nl/monica/Cell type-specific-complex-formation-of-key-transcription-factors-in-the-root-SCN) and in GitHub (https://github.com/moneralee/Cell type-specific-complex-formation-of-key-transcription-factors-in-the-root-SCN) upon publication.

## Acknowledgements

We would like to acknowledge funding of V.I.S. by the Deutsche Forschungsgesellschaft (DFG) through grant STA1212/4-1 to Y.S. M.L.G.G. is supported by the long-term program PlantXR: A new generation of breeding tools for extra-resilient crops (KICH3.LTP.20.005) which is financed by the Dutch Research Council (NWO), the Foundation for Food & Agriculture Research (FFAR), companies in the plant breeding and processing industry, and Dutch universities. These parties collaborate in the CropXR Institute (www.cropxr.org) that is funded through the National Growth Fund (NGF) of the Netherlands. We thank Rebecca C. Burkart for sharing PLT3, PLT3ΔPrD and WOX5 constructs and stable Arabidopsis lines. We thank Ana Can᷉o-Delgado for sharing seeds of *bravo-2* single and *bravo-2 wox5-1* double mutants. We thank Kirsten ten Tusscher for insightful discussions on the modelling and Jan Kees van Amerongen for management of computational facilities of the Theoretical Biology group (Utrecht University). We thank Cornelia Gieseler, Carin Theres and Silke Winters for technical assistance. We thank Meik H. Thiele for help with statistical analyses and data visualisation with R. We also thank Jan E. Maika for help with fitting FRET-FLIM data. We would like to acknowledge the Center for Advanced Imaging (CAi) at Heinrich-Heine-University Düsseldorf for providing access to the Zeiss LSM780 and LSM880 and especially Dr. Sebastian Hänsch and Prof. Dr. Stefanie Weidtkamp-Peters for general support during imaging and analysis. Funding for instrumentation: Zeiss LSM 780: DFG-INST 208/551-1 FUGG and Zeiss LSM 880: DFG-INST 208/746-1 FUGG.

## Author contributions

Y.S. conceived the project. Y.S. and V.I.S. designed, analyzed and interpreted the data. V.I.S. carried out all experiments. M.L.G.G. formulated and performed mathematical modelling. The manuscript was written by V.S. and M.L.G.G. and was revised by Y.S. All authors commented and approved the manuscript.

## Declaration of competing interests

The authors declare no competing interests.

This manuscript has not been accepted or published elsewhere.

## Notes

### Competing Interest Statement

The authors have declared no competing interest.

