## Supplemental data for "Stem cell homeostasis in the root of *Arabidopsis* involves cell type specific complex formation of key transcription factors"

### Appendix

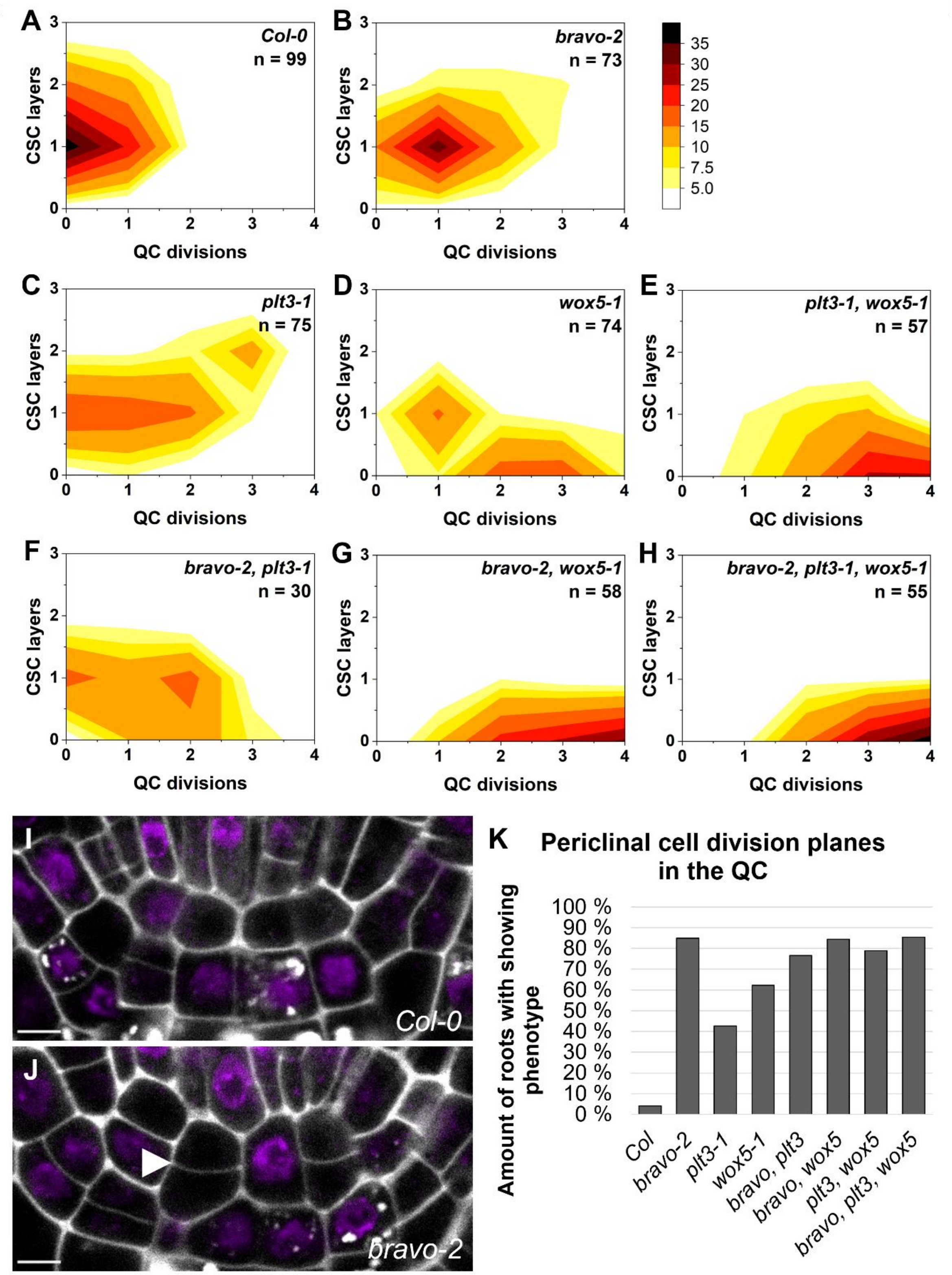

**Figure S1: Elevated QC division frequencies negatively correlate to the number of CSC layers. A-H.** 2D histograms visualizing the combined results of the SCN staining in the respective genotype showing the number of CSC layers on the y-axis and QC divisions on the x-axis. Darker colours correspond to a higher number of roots showing the phenotype. Number of analysed roots per genotype (biological replicate) is indicated in each graph and results from 3-5 technical replicates. **I.** Close-up of the QC in the Col-0 WT and **J.** in the *bravo-2* mutant showing an additional periclinal cell division plane (white arrowhead). **K.** Quantification of periclinal cell division planes in the different mutants.

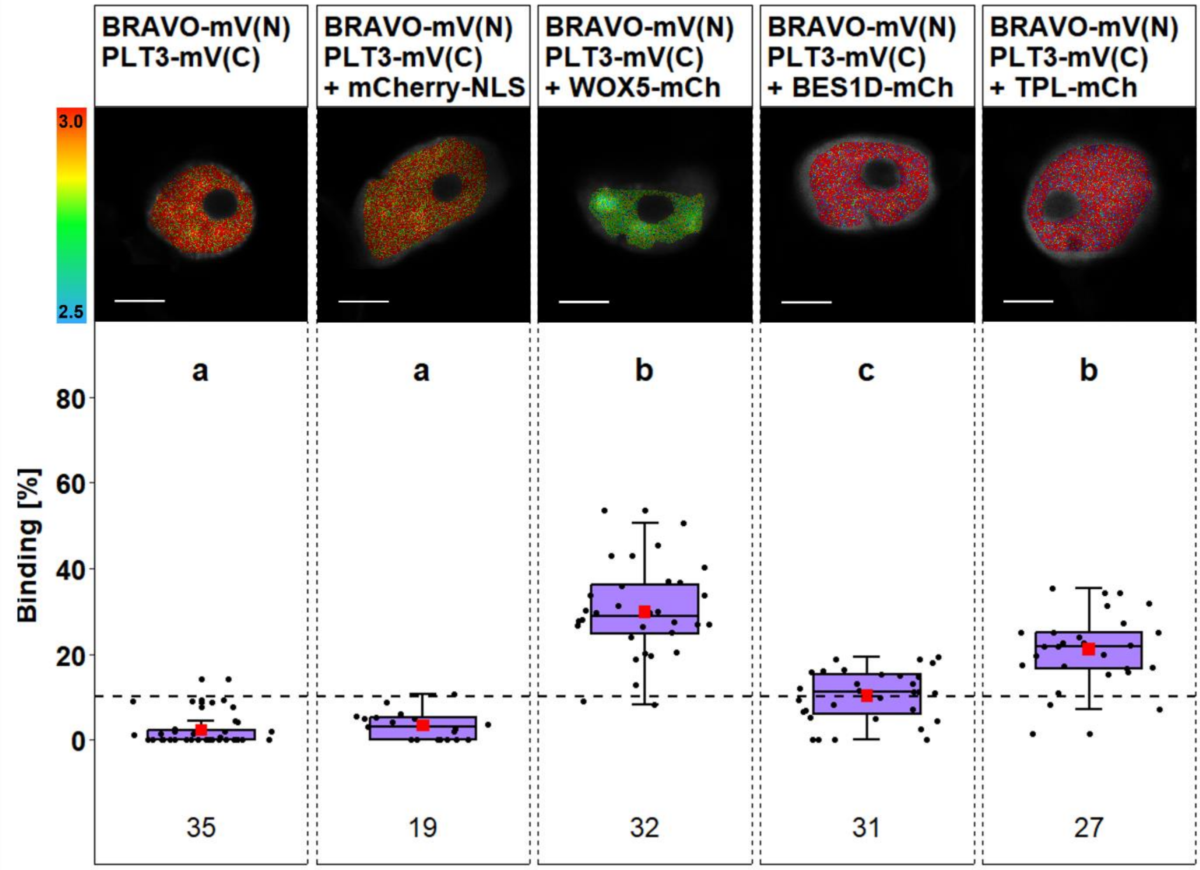

**Figure S2: Trimeric complex formation of BRAVO and PLT3 with WOX5, BES1D and TPL. Upper panel**: Representative images of fluorescence lifetime imaging microscopy (FLIM) measurements in *N. benthamiana* epidermal leaf cells after a pixel-wise mono- or biexponential fit. The fluorescence lifetime of the donor BRAVO-mV(N) PLT3-mV(C) in presence or absence of the indicated acceptor is color-coded: blue (2.5) refers to low fluorescence lifetime [in ns], red (3.0) indicates high fluorescence lifetime. Scale bar represents 6 µm. **Lower panel:** Binding [%] (magenta) for BRAVO-mV(N) PLT3-mV(C) with or without co-expression of mCherry-NLS, WOX5-mCh, BES1D-mCh or TPL-mCh. Statistical groups were assigned after a non-parametric Kruskal Wallis ANOVA with *post-hoc* Dunn's test (α = 0.05). Black dotted line indicates the Binding cut-off of 10 %. Number of analysed nuclei is indicated below each sample and results from 3-4 technical replicates.

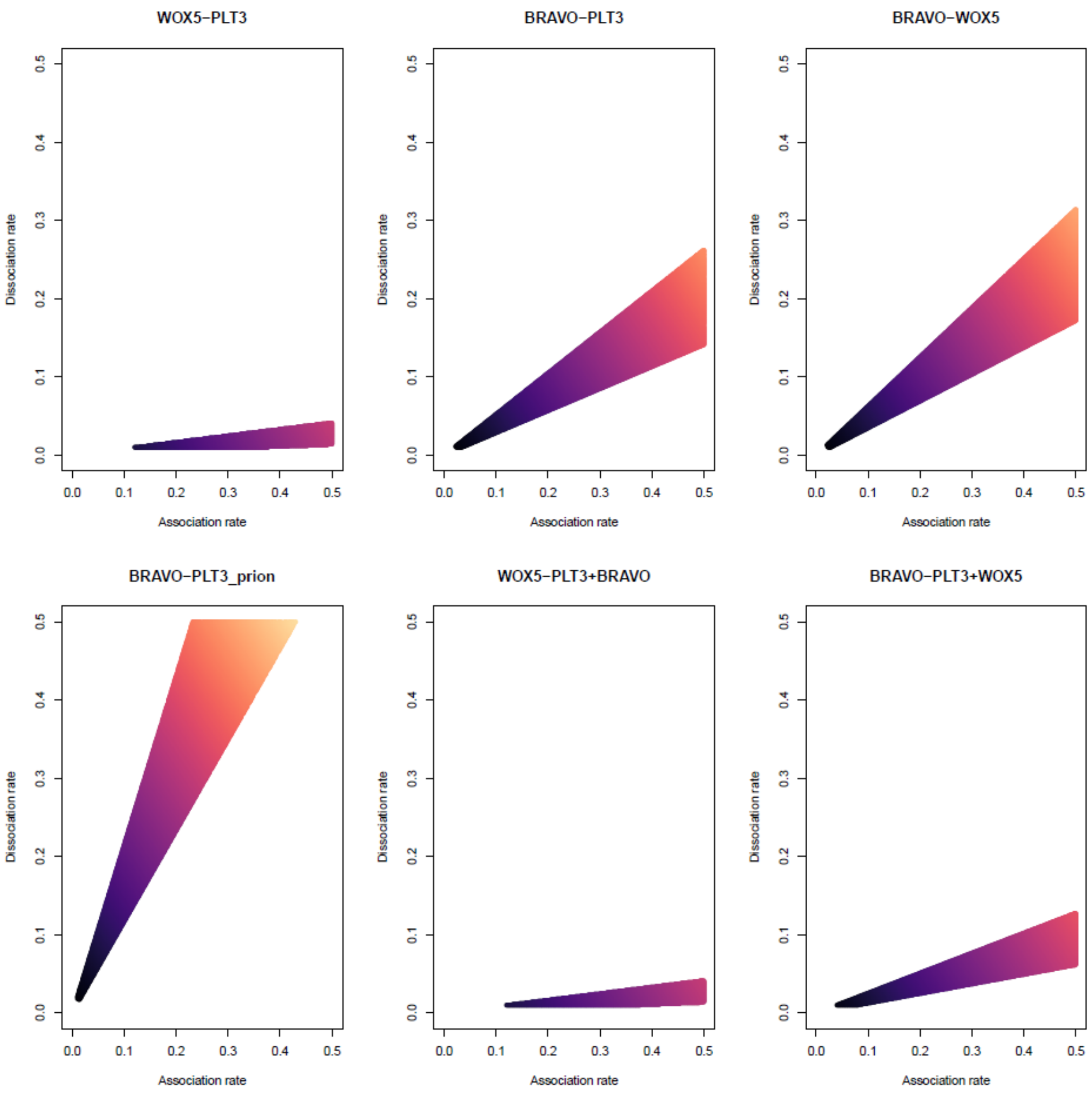

**Figure S3: Association and dissociation parameters predicted for the heterodimers and trimeric complex modelled.** For each protein complex studied we show the combination of association and dissociation parameters (colored area) that can produce the protein complex formation in agreement with the binding affinities described experimentally.

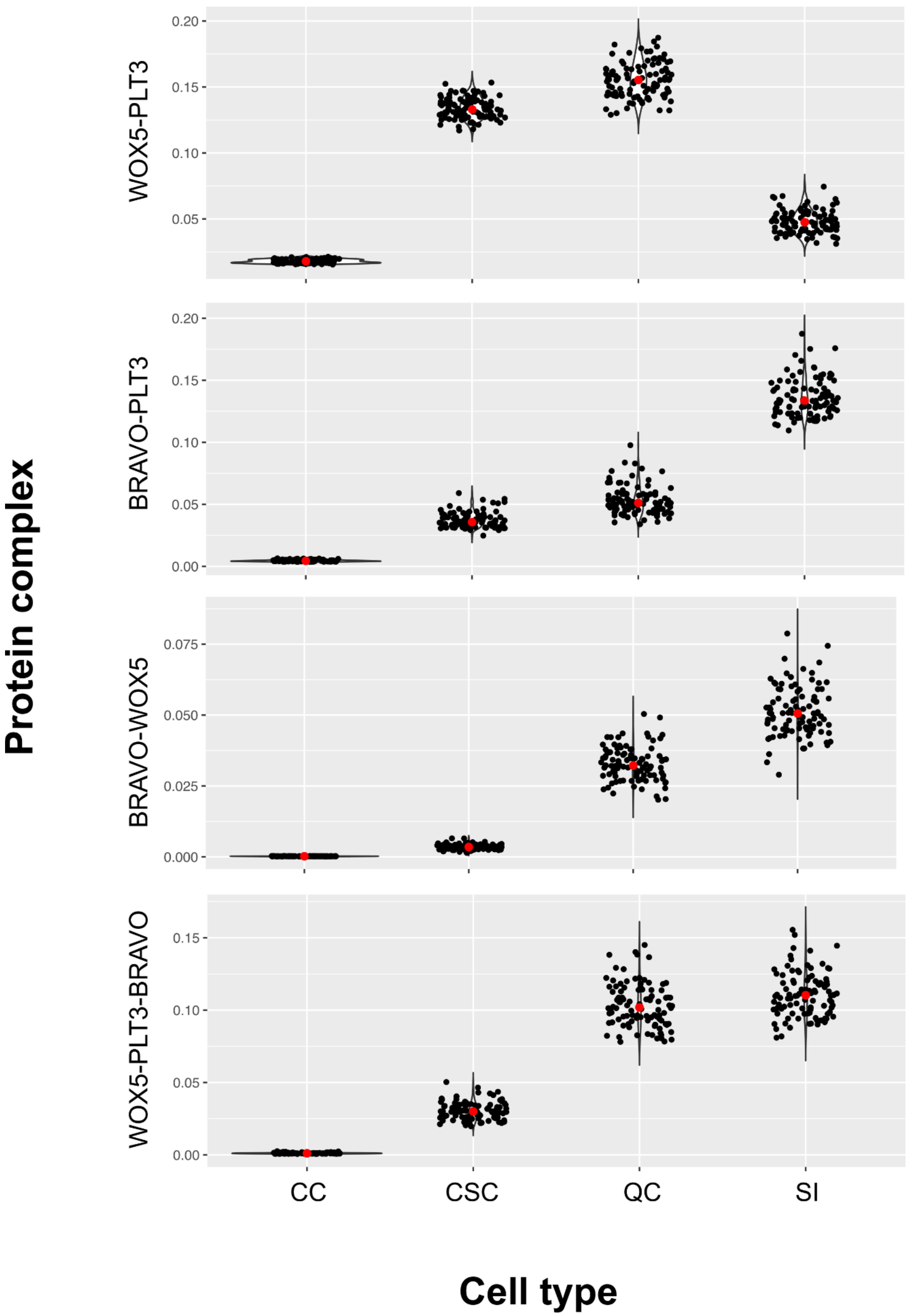

**Figure S4: Robustness of the protein complex cell signatures.** Simulations of the formation of protein complexes in the cells of the root stem cell niche using 100 different association and dissociation parameter sets. The resulting levels of each simulation for WOX5-PLT3, BRAVO-PLT3, BRAVO-WOX5 and WOX5-PLT3-BRAVO in each cell type are shown, indicating they are robust to the specific parameters used in the simulations. Mean values shown in red.

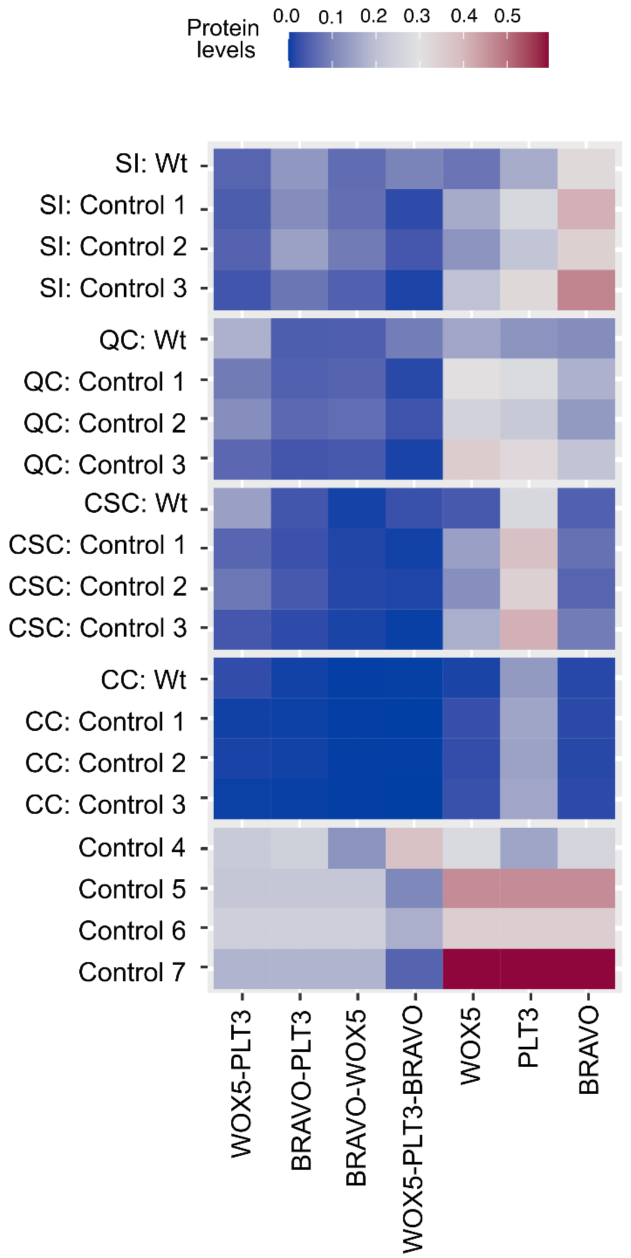

**Figure S5:Controls for *in silico* prediction of protein complex signatures in the WT root SCN.** For Control 1-3, the experimentally determined protein abundances were used, combined with the assumptions that association and dissociation rates are equal, a higher association and a higher dissociation rate, respectively. For Control 4, experimentally determined association and dissociation rates were used in combination with equal protein abundances among all cell-types and TFs. Controls 5-7 combine equal protein levels with assumed association/dissociation rates from Control 1-3, respectively. Heatmap showing the protein complexes and free protein in the cells of WT simulation. High concentrations are displayed in red, low concentration are displayed in blue. SI: stele initals; QC: quiescent center; CSC: columella stem cells; CC: columella cells.

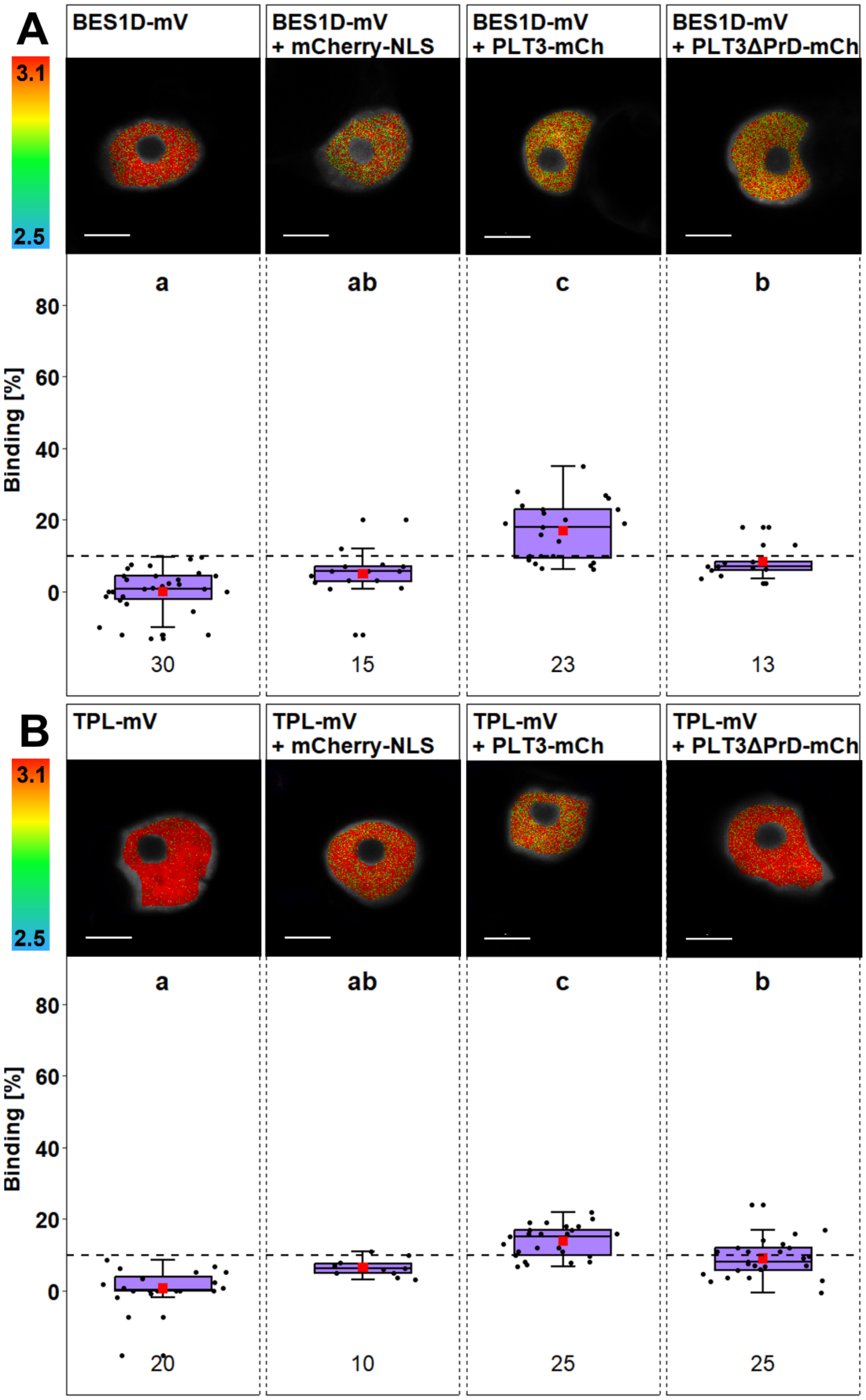

**Figure S6: Interaction of PLT3 with BES1D and TPL depends on PrDs found in PLT3. Upper panels:** Representative images of fluorescence lifetime imaging microscopy (FLIM) measurements in *N. benthamiana* epidermal leaf cells after a pixel-wise mono- or biexponential fit. The fluorescence lifetime of the donor BES1DmV (A) or TPL-mV (B) in presence or absence of the indicated acceptor is color-coded: blue (2.5) refers to low fluorescence lifetime [in ns], red (3.1) indicates high fluorescence lifetime. Scale bar represents 6 µm. Lower panels: Binding [%] (magenta) for BES1D-mV (A) or TPL-mV (B) with or without co-expression of mCherry-NLS, PLT3-mCh or PLT3dPrD-mCh. Statistical groups were assigned separately for experiments with BES1D-mV and TPL-mV after a non-parametric Kruskal Wallis ANOVA with *post-hoc* Dunn's test ($\alpha = 0.05$). Black dotted line indicates the Binding cut-off of 10 %. Number of analysed nuclei is indicated below each sample and results from 2-3 technical replicates.

**Table S1:** List of primers used for cloning. Italic bases represent overhangs and *Bsa*I recognition sites necessary for GreenGate cloning.

| Gene identifier | alias | Primer name | Orientation | Sequence 5'-3' orientation |
|---|---|---|---|---|
| Promoter modules | | | | |
| AT5G17800 | BRAVO | VS_GG_pBRAVO_F | F | *AAAGGTCTCAACCT*CCACTAACCATTTCGTAA |
| | | VS_GG_pBRAVO_R | R | *AAAGGTCTCATGTT*GTTTCTGGTTTAGGGATTA |
| CDS in C modules | | | | |
| AT1G19350 | BES1 | VS_GG_BES1D_CDS_F | F | *AAAGGTCTCAGGCTTA*ATGACGTCTGACGGAGCAAC |
| | | VS_GG_BES1D_CDS_R | R | *AAAGGTCTCACTGA*ACTATGAGCTTTACCATTTCC |
| AT5G17800 | BRAVO | RD_GG_BRAVO F | F | *AAAGGTCTCAGGCTTA*ATGAATCCAAATC |
| | | RD_GG_BRAVO R | R | *AAAGGTCTCACTGA*GGAAGCTCCAAC |
| AT1G15750 | TPL | VS_GG_TPL_CDS_F | F | *AAAGGTCTCAGGCTTA*ATGTCTTCTCTTAG |
| | | VS_GG_TPL_CDS_R | F | *AAAGGTCTCACTGA*TCTCTGAGGCTG |

**Table S2:** List of primers used for genotyping.

| Gene ID | SALK ID | alias | Primer | | |
|---|---|---|---|---|---|
| | | | Orientation | Name | Sequence 5'-3' |
| AT5G17800 | SALK_062413 | *bravo-2* | F | VS_bravo-2_F | TCCCTTAATCCCTAAACCCAGC |
| | | | R | VS_bravo-2_R | CCTGATGCAAGGGTACTATCG |
| AT3G11260 | SALK_038262 | *wox5-1* | F | GK_WOX5 F | AAACAGTTGAGGACTTTACATCTGA |
| | | | R | WOX5 R | CGGATAATATGTCATAATTCAAAAT |
| AT5G10510 | SALK_127417 | *plt3-1* | F | GK_PLT3L | TTGTGATTTGCCATTGACTAAAGGT |
| | | | R | GK_PLT3R | GAAAACAGTCCAATGGTCTCACATC |

**Table S3:** List of entry vectors used for GreenGate cloning.

| Name | Module (Backbone) | Insert | Reference |
|---|---|---|---|
| pGGB002 | B | Omega element | (Lampropoulos et al. 2013) |
| pGGD007 | D | Linker-NLS | |
| pGGE009 | E | UBIQUITIN 10 terminator | |
| pGGF002 | F | BASTA resistance | |
| pGGG001 | G | Adapter | |
| pGGG002 | G | Adapter | |
| pGGM000 | M | Empty intermediate vector | |
| pGGN000 | N | Empty intermediate vector | |
| pGGZ001 | Z | Empty destination vector | |
| pRD42 | C | mVenus | (Burkart et al. 2022) |
| pRD43 | D | mVenus | |
| pRD53 | D | mCherry | |
| pRD45 | A | *WOX5* promoter | |
| pRD40 | C | WOX5 CDS | |
| pRD41 | C | PLT3 CDS | |
| pRD65 | B | Glucocorticoid receptor | |
| pRD101 | C | PLT3dPrD | |
| pPD161 | A | Ubi-XVE oLexA-35S | (Denninger et al. 2019) |
| pVS125 | A | *BRAVO* promoter | This study |
| pRD135 | C | BRAVO CDS | |
| pVS191 | C | BES1D | |
| pVS84 | C | TPL | |
| pJM81 | D | mVenus(N) | (Maika et al. 2023) |
| pJM82 | D | mVenus(C) | |
| pBLAD011 | D | mTurquoise2 | |

**Table S4:** List of expression vectors for stable transformation of *A. thaliana* or transient transformation of *N. benthamiana* generated in this study.

| Plasmid ID | Construct | GreenGate module | | | | | | | Resistance |
|---|---|---|---|---|---|---|---|---|---|
| | | A | B | C | D | E | F | Z | |
| pVS133 | pBRAVO:BRAVO-mV | *BRAVO* promoter | Ω element (pGGB002) | BRAVO | mVenus | UBQ10 terminator (pGGE009) | BASTA R (pGGF002) | pGGZ001 | Spec |
| pVS139 | Inducible BES1D-mVenus | Ubi-XVE oLexA-35S | Ω element (pGGB002) | BES1D | mVenus | UBQ10 terminator (pGGE009) | BASTA R (pGGF002) | pGGZ001 | Spec |
| pVS140 | Inducible BES1D-mCherry | Ubi-XVE oLexA-35S | Ω element (pGGB002) | BES1D | mCherry | UBQ10 terminator (pGGE009) | BASTA R (pGGF002) | pGGZ001 | Spec |
| pVS141 | Inducible BRAVO-mVenus | Ubi-XVE oLexA-35S | Ω element (pGGB002) | BRAVO | mVenus | UBQ10 terminator (pGGE009) | BASTA R (pGGF002) | pGGZ001 | Spec |
| pVS142 | Inducible BRAVO-mCherry | Ubi-XVE oLexA-35S | Ω element (pGGB002) | BRAVO | mCherry | UBQ10 terminator (pGGE009) | BASTA R (pGGF002) | pGGZ001 | Spec |
| pVS143 | Inducible TPL-mCherry | Ubi-XVE oLexA-35S | Ω element (pGGB002) | TPL | mCherry | UBQ10 terminator (pGGE009) | BASTA R (pGGF002) | pGGZ001 | Spec |
| pVS85 | Inducible TPL-mVenus | Ubi-XVE oLexA-35S | Ω element (pGGB002) | TPL | mVenus | UBQ10 terminator (pGGE009) | BASTA R (pGGF002) | pGGZ001 | Spec |
| pVS154 | Inducible WOX5-mVenus(N) | Ubi-XVE oLexA-35S | Ω element (pGGB002) | WOX5 | mVenus(N) | UBQ10 terminator (pGGE009) | - | pGGM000 | Kan |
| pVS156 | Inducible PLT3-mVenus(C) | Ubi-XVE oLexA-35S | Ω element (pGGB002) | PLT3 | mVenus(C) | UBQ10 terminator (pGGE009) | BASTA R (pGGF002) | pGGN000 | Kan |
| pVS163 | Inducible WOX5-mVenus(N)/ Inducible PLT3-mVenus(C) | pVS154 + pVS156 | | | | | | pGGZ001 | Spec |
| pVS167 | Inducible nuclear localized mCherry | Ubi-XVE oLexA-35S | Ω element (pGGB002) | mCherry (pGGC015) | linker-NLS (pGGD007) | UBQ10 terminator (pGGE009) | BASTA R (pGGF002) | pGGZ001 | Spec |
| pVS180 | Inducible BRAVO-mVenus(N) | Ubi-XVE oLexA-35S | Ω element (pGGB002) | BRAVO | mVenus(N) | UBQ10 terminator (pGGE009) | - | pGGM000 | Kan |

| | | | | | | | | | |
|---|---|---|---|---|---|---|---|---|---|
| pVS233 | Inducible BRAVO-mVenus(N)/ Inducible PLT3-mVenus(C) | pVS180 + pVS156 | | | | | | pGGZ001 | Kan |
| pVS288 | pWOX5:GR-PLT3-mTurquoise2 | *WOX5* promoter | GR | PLT3 | mT2 | UBQ10 terminator (pGGE009) | BASTA R (pGGF002) | pGGZ001 | Spec |
| pVS289 | pWOX5:GR-PLT3ΔPrD-mTurquoise2 | *WOX5* promoter | GR | PLT3ΔPrD | mT2 | UBQ10 terminator (pGGE009) | BASTA R (pGGF002) | pGGZ001 | Spec |

**Table S5:** List of Arabidopsis mutants and transgenic lines used in this study.

| Gene ID | Alias | Reference |
|---|---|---|
| AT5G17800 | *bravo-2* | (Vilarrasa-Blasi et al. 2014) |
| | *Col-0, pBRAVO:BRAVO-mVenus* | This study by dipping |
| AT5G10510 | *plt3-1* | (Galinha et al. 2007) |
| | *Col-0, pPLT3:PLT3-mVenus* | (Burkart et al. 2022) |
| AT5G17800, AT5G10510 | *bravo-2, plt3-1* | This study by crossing of *bravo-2* and *plt3-1* |
| AT3G11260 | *wox5-1* | (Burkart et al. 2022) |
| | *Col-0, pWOX5:WOX5-mVenus* | |
| AT5G10510, AT3G11260 | *plt3-1, wox5-1* | (Burkart et al. 2022) |
| | *plt3-1, pWOX5:GR-PLT3-mTurquoise2* | This study by dipping |
| | *plt3-1, pWOX5:GR-PLT3ΔPrD-mTurquoise2* | This study by dipping |
| AT5G17800, AT3G11260 | *bravo-2, wox5-1* | (Betegón-Putze et al. 2021) |
| AT5G17800, AT5G10510, AT3G11260 | *bravo-2, plt3-1, wox5-1* | This study by crossing of *bravo-2* and *plt3-1, wox5-1* |

**Table S6:** Fluorescence intensities of *pPLT3:PLT3-mV, pBRAVO:BRAVO-mV* and *pWOX5:WOX5-mV* translational reporter in different cell types corresponding to Fig. 1, 5 and 7.

| Fluorescence intensity | Date | Cell type | | | |
|---|---|---|---|---|---|
| | | SI | QC | CSC | CC |
| *pPLT3:PLT3-mV* | 28.11.23 | 30360.17 | 37523.45 | 48567.14 | 17482.71 |
| | | 19709.9 | 18540.11 | 22274.4 | 15590.86 |
| | | 19782.11 | 16327.93 | 26233.66 | 6335.124 |
| | | 24665.88 | 22914.56 | 22039.49 | 13233.82 |
| | | 16336.64 | 14473.61 | 16777.05 | 7263.823 |
| | | 20659.51 | 17532.76 | 19131.18 | 9199.955 |
| | | 48822.24 | 37769.87 | 55574.26 | 5490.301 |
| | | 42071.03 | 23881.81 | 28888.19 | 6474.87 |
| | | 31306.43 | 28682.1 | 43367.48 | 26115.04 |
| | | 26035.98 | 24754.96 | 37376.88 | 5407.617 |
| | AV | 27974.99 | 24240.11 | 32022.97 | 11259.41 |
| | SD | 9945.883 | 7853.973 | 12737.42 | 6457.178 |
| | 08.12.23 | 13458.53 | 12022.34 | 10693.72 | 3079.556 |
| | | 11382.8 | 19661.37 | 23237.77 | 2543.246 |
| | | 31222.01 | 28196.38 | 24778.53 | 1576.29 |
| | | 28921.69 | 24819.56 | 21161.82 | 6278.363 |
| | | 25231.6 | 12692.34 | 9225.459 | 9485.656 |
| | | 14733.96 | 15308.52 | 30900.44 | 19996.52 |
| | | 13009.25 | 19128.18 | 18887.84 | 1805.671 |
| | | 17399.32 | 20006.95 | 19149.94 | 6983.197 |
| | | 10872.27 | 8227.07 | 12223.45 | 1741.223 |
| | | 21828.1 | 21030.97 | 24277.79 | 2664.413 |
| | AV | 18805.95 | 18109.37 | 19453.67 | 5615.413 |
| | SD | 7111.341 | 5781.51 | 6587.363 | 5426.973 |
| | 28.12.23 | 36705.69 | 30759.92 | 32179.98 | 13431.11 |
| | | 23201.13 | 22910.14 | 26746.3 | 12992.69 |
| | | 14505.16 | 16790.7 | 14650.75 | 10778.34 |
| | | 25326.28 | 16618.53 | 15902.32 | 7739.259 |
| | | 22976.1 | 22252.79 | 13183.53 | 15139.32 |
| | | 26289.13 | 34416.08 | 46981.29 | 13944.07 |
| | | 23456.61 | 45225.47 | 45902.98 | 7467.853 |
| | | 19359.14 | 19409.83 | 23114.97 | 23172.93 |
| | AV | 23977.41 | 26047.93 | 27332.76 | 13083.19 |
| | SD | 5938.235 | 9373.698 | 12570.98 | 4640.888 |
| | Overall AV | 23267.05 | 22671.05 | 25904.29 | 9224.714 |
| | Overall SD | 8745.45 | 8164.877 | 11578.14 | 6400.249 |
| *pWOX5:WOX5-mV* | 28.11.23 | 29201.12 | 45834.2 | 10795.79 | 896.549 |
| | | 13869.69 | 21386.9 | 8486.974 | 1496.997 |
| | | 19027.65 | 28070.95 | 7576.267 | 779.7283 |
| | | 14906.16 | 27262.89 | 9271.191 | 685.0723 |
| | | 15691.41 | 27641.05 | 6953.646 | 687.679 |
| | | 17316.21 | 20991.05 | 10378.01 | 705.1347 |

| | | | | | |
|---|---|---|---|---|---|
| | | 11098.75 | 30414.8 | 9765.741 | 766.4405 |
| | | 29355.06 | 22623.3 | 7448.115 | 882.8665 |
| | | 18739.97 | 22616.32 | 9814.987 | 1002.133 |
| | | 13616.57 | 17752.29 | 12525.81 | 7864.905 |
| | **AV** | 18282.26 | 26459.37 | 9301.653 | 1576.75 |
| | **SD** | 5953.816 | 7460.556 | 1638.761 | 2108.523 |
| | **08.12.23** | 11408.16 | 12757.81 | 7483.419 | 532.5793 |
| | | 14144.85 | 17938.5 | 8482.929 | 958.597 |
| | | 18676.1 | 20097.02 | 12921.63 | 838.317 |
| | | 18215.11 | 22406.33 | 10450.67 | 730.0777 |
| | | 12329.53 | 17780.14 | 8244.796 | 3459.093 |
| | | 32552.77 | 32325.28 | 15897.64 | 8156.247 |
| | | 18057.62 | 34627.05 | 17197.87 | 833.5587 |
| | | 20081.02 | 33351.82 | 12183.4 | 2112.58 |
| | | 16962.96 | 27887.07 | 13940.99 | 854.592 |
| | | 11985.38 | 14764.86 | 6683.364 | 731.142 |
| | **AV** | 17441.35 | 23393.59 | 11348.67 | 1920.678 |
| | **SD** | 5841.4 | 7655.123 | 3466.346 | 2246.346 |
| | **28.12.23** | 18809.39 | 21126.97 | 13228.5 | 969.778 |
| | | 18091.15 | 21354.1 | 13364.58 | 661.5485 |
| | | 13782.29 | 25015.66 | 7060.959 | 837.938 |
| | | 11749.74 | 32733.76 | 10815.74 | 2313.319 |
| | | 16980.22 | 19147.65 | 10490.34 | 719.408 |
| | | 16397.84 | 28137.24 | 15536.71 | 1107.079 |
| | | 17359.42 | 26238.14 | 10937.61 | 680.671 |
| | | 16533.92 | 36878.85 | 15549.16 | 7561.397 |
| | **AV** | 16213 | 26329.05 | 12122.95 | 1856.392 |
| | **SD** | 2183.395 | 5708.609 | 2682.338 | 2214.819 |
| | **Overall AV** | 16790.78 | 25132.08 | 11061.31 | 1684.39 |
| | **Overall SD** | 5507.95 | 7039.065 | 3596.461 | 2105.804 |
| ***pBRAVO:BRAVO-mV*** | **28.11.23** | 33296 | 16940.47 | 4615.491 | 852.569 |
| | | 32465.25 | 12767.84 | 8112.653 | 989.9733 |
| | | 36508.05 | 19942.11 | 14826.77 | 1133.772 |
| | | 31225.91 | 18182.72 | 8115.524 | 919.5017 |
| | | 40549.32 | 20651.28 | 8776.193 | 1081.686 |
| | | 40143.53 | 29515.55 | 8153.343 | 1102.827 |
| | | 19431.36 | 10567.79 | 3988.657 | 695.031 |
| | | 29660.65 | 12654.5 | 4704.077 | 1479.854 |
| | | 28870.52 | 16931.86 | 3053.96 | 872.8765 |
| | | 31988.38 | 16361 | 8163.813 | 1257.115 |
| | **AV** | 32413.9 | 17451.51 | 7251.048 | 1038.52 |
| | **SD** | 5777.106 | 5065.289 | 3233.365 | 213.1088 |
| | **08.12.23** | 17139.23 | 12128.37 | 3986.173 | 845.2763 |
| | | 35034.8 | 21779.82 | 10615.84 | 1246.433 |
| | | 32923.36 | 12882.26 | 6763.008 | 946.799 |
| | | 19276.64 | 14514.07 | 9689.346 | 906.56 |
| | | 25266.62 | 12037.34 | 6404.131 | 734.922 |

| | | | | | |
|---|---|---|---|---|---|
| | | 41025.65 | 18926.21 | 7023.33 | 1303.169 |
| | | 24780.18 | 10123.88 | 3126.593 | 844.1405 |
| | | 39899.04 | 18246.41 | 10925.48 | 1296.336 |
| | | 22975.67 | 13452.88 | 11498.59 | 1244.351 |
| | | 31024.35 | 13617 | 9103.976 | 1312.867 |
| | **AV** | 28934.55 | 14770.82 | 7913.646 | 1068.085 |
| | **SD** | 7890.066 | 3481.702 | 2766.423 | 219.4884 |
| | **28.12.23** | 46147.85 | 26669.37 | 16433.59 | 1442.835 |
| | | 49948.21 | 7967.577 | 2823.43 | 770.814 |
| | | 39406.01 | 34224.94 | 6801.504 | 1497.628 |
| | | 41437.48 | 16972.48 | 5323.483 | 1414.79 |
| | | 25584.47 | 9576.138 | 2069.954 | 805.6805 |
| | | 40467.97 | 9512.593 | 5478.113 | 1207.251 |
| | | 42555.79 | 10385.78 | 5205.178 | 1349.889 |
| | | 30363.89 | 13326.26 | 9106.051 | 1152.85 |
| | **AV** | 39488.96 | 16079.39 | 6655.162 | 1205.217 |
| | **SD** | 7454.075 | 8898.174 | 4222.165 | 264.0782 |
| | **Overall AV** | 33047.28 | 15892.74 | 6833.518 | 1058.715 |
| | **Overall SD** | 7897.687 | 6132.839 | 3588.269 | 257.7053 |

AV: average, STD: standard deviation.

**Table S7:** Average number of QC divisions and CSC layers per root related to Fig. 2 and S1.

| Genotype | Average number of QC divisions per root | Average number of CSC layers per root | Number of analysed roots |
|---|---|---|---|
| *Col-0* | 0.535354 | 1.282828 | 99 |
| *bravo-2* | 1.30137 | 1.123288 | 73 |
| *plt3-1* | 1.386667 | 1.293333 | 75 |
| *wox5-1* | 1.90541 | 0.594595 | 74 |
| *bravo, plt3* | 1.5333333 | 0.73333333 | 30 |
| *bravo, wox5* | 2.844827586 | 0.103448276 | 58 |
| *plt3, wox5* | 2.7719298 | 0.2631579 | 57 |
| *bravo, plt3, wox5* | 3.145454545 | 0.163636364 | 55 |

**Table S8:** Ratio of periclinal cell division planes in the QC related to Fig. S1.

| Genotype | Periclinal cell division planes in the QC [%] | Number of analysed roots |
|---|---|---|
| *Col-0* | 4 | 99 |
| *bravo-2* | 85 | 73 |
| *plt3-1* | 43 | 75 |
| *wox5-1* | 62 | 78 |
| *bravo, plt3* | 77 | 30 |
| *bravo, wox5* | 84 | 59 |
| *plt3, wox5* | 79 | 57 |
| *bravo, plt3, wox5* | 85 | 55 |

**Table S9:** Measured FRET efficiency and Binding values related to Fig. 3, 5 and 7.

| **Sample** | **BRAVO-mV** | | **BRAVO-mV mCherry-NLS** | | **BRAVO-mV PLT3-mCh** | | **BRAVO-mV BES1D-mCh** | | **BRAVO-mV WOX5-mCh** | | **BRAVO-mV TPL-mCh** | |
|---|---|---|---|---|---|---|---|---|---|---|---|---|
| **Date** | **FRET E** | **Binding** | **FRET E** | **Binding** | **FRET E** | **Binding** | **FRET E** | **Binding** | **FRET E** | **Binding** | **FRET E** | **Binding** |
| **20.12.19** | 80.00 | 3.00 | | | 54.00 | 34.00 | | | | | | |
| | 80.00 | 1.70 | | | 51.00 | 33.00 | | | | | | |
| | 80.00 | 2.40 | | | 56.00 | 35.00 | | | | | | |
| | 80.00 | 7.10 | | | 54.00 | 30.00 | | | | | | |
| | 80.00 | 5.30 | | | 60.00 | 40.00 | | | | | | |
| | 80.00 | 2.50 | | | 53.00 | 45.00 | | | | | | |
| | 80.00 | 8.00 | | | 57.00 | 35.00 | | | | | | |
| | 80.00 | 8.90 | | | 53.00 | 40.00 | | | | | | |
| | 80.00 | 5.10 | | | 58.00 | 32.00 | | | | | | |
| | 78.00 | 8.00 | | | 59.00 | 34.00 | | | | | | |
| **04.02.20** | 9.88 | -11.00 | | | | | 80.00 | 9.20 | | | | |
| | 9.88 | -17.00 | | | | | 0.97 | 7.00 | | | | |
| | 17.00 | 7.40 | | | | | 0.66 | 37.00 | | | | |
| | 10.00 | -0.30 | | | | | 0.90 | 13.00 | | | | |
| | 71.00 | 6.30 | | | | | 0.49 | 34.00 | | | | |
| | 80.00 | 6.90 | | | | | 0.93 | 15.00 | | | | |
| | 80.00 | 8.20 | | | | | 1.10 | 22.00 | | | | |
| | 9.88 | -23.00 | | | | | 0.64 | 40.00 | | | | |
| | 67.00 | 4.30 | | | | | 1.30 | 9.05 | | | | |
| | 16.00 | 5.00 | | | | | | | | | | |
| **17.02.20** | 80.00 | 2.50 | 45.00 | 11.00 | | | 48.00 | 29.00 | 58.00 | 9.60 | 43.00 | 24.00 |
| | 80.00 | 0.51 | 43.00 | 8.41 | | | 48.00 | 25.00 | 55.00 | 27.00 | 39.00 | 27.00 |
| | 41.00 | 8.20 | 63.00 | 8.90 | | | 46.00 | 35.00 | 64.00 | 14.00 | 37.00 | 30.00 |
| | 80.00 | 2.40 | 50.00 | 15.00 | | | 47.00 | 19.00 | 61.00 | 14.00 | 46.00 | 28.00 |
| | 10.10 | -11.00 | 53.00 | 14.00 | | | 41.00 | 29.00 | 56.00 | 33.00 | 40.00 | 29.00 |
| | 75.00 | 2.20 | | | | | 42.00 | 30.00 | 58.00 | 19.00 | 43.00 | 30.00 |
| | 80.00 | 1.00 | | | | | 45.00 | 33.00 | 59.00 | 27.00 | 40.00 | 34.00 |
| | 80.00 | 4.10 | | | | | 45.00 | 47.00 | 51.00 | 7.30 | 42.00 | 32.00 |

| | | | | | | | | | | | | |
|---|---|---|---|---|---|---|---|---|---|---|---|---|
| | 80.00 | 1.40 | | | | | 45.00 | 41.00 | 64.00 | 7.00 | 49.00 | 20.00 |
| | 80.00 | 6.70 | | | | | 48.00 | 49.00 | 65.00 | 11.00 | 38.00 | 29.00 |
| | | | | | | | 41.00 | 36.00 | 40.00 | 24.00 | 48.00 | 27.00 |
| | | | | | | | 45.00 | 41.00 | 49.00 | 27.00 | 44.00 | 32.00 |
| | | | | | | | 48.00 | 42.00 | 52.00 | 31.00 | 77.00 | 73.00 |
| | | | | | | | 45.00 | 37.00 | 50.00 | 39.00 | 43.00 | 37.00 |
| | | | | | | | 44.00 | 39.00 | 55.00 | 61.00 | 43.00 | 34.00 |
| **17.08.20** | 80.00 | 6.40 | 80.00 | 12.00 | 74.00 | 8.70 | | | 79.00 | 15.00 | 59.00 | 16.00 |
| | 80.00 | 8.20 | 66.00 | 9.60 | 61.00 | 28.00 | | | 66.00 | 10.00 | 57.00 | 18.00 |
| | 80.00 | 11.00 | 68.00 | 11.00 | 80.00 | 10.00 | | | 65.00 | 17.00 | 55.00 | 23.00 |
| | 80.00 | 11.00 | 80.00 | 8.90 | 80.00 | 8.70 | | | 56.00 | 24.00 | 56.00 | 22.00 |
| | 80.00 | 14.00 | 80.00 | 13.00 | 73.00 | 10.70 | | | 55.00 | 26.00 | 56.00 | 20.00 |
| | 80.00 | 5.50 | 77.00 | 8.70 | 77.00 | 12.00 | | | 63.00 | 14.00 | 50.00 | 26.00 |
| | 80.00 | 6.40 | 80.00 | 12.00 | 80.00 | 9.60 | | | | | 76.00 | 10.00 |
| | 80.00 | 3.50 | 78.00 | 15.00 | 62.00 | 16.00 | | | | | 48.00 | 25.00 |
| | 80.00 | 6.50 | | | 55.00 | 24.00 | | | | | 53.00 | 20.80 |
| | 80.00 | 3.20 | | | 52.00 | 25.00 | | | | | 52.00 | 23.00 |
| **27.06.22** | 35.00 | 1.10 | 63.00 | 1.40 | 49.00 | 14.00 | 47.00 | 13.00 | 44.00 | 13.00 | 44.00 | 17.00 |
| | 80.00 | 3.50 | 80.00 | 3.90 | 45.00 | 40.00 | 45.00 | 21.00 | 59.00 | 9.10 | 43.00 | 25.00 |
| | 78.00 | 5.00 | 63.00 | 2.40 | 43.00 | 26.00 | 44.00 | 19.00 | 43.00 | 18.00 | 38.00 | 29.00 |
| | 80.00 | -0.15 | 54.00 | 2.10 | 48.00 | 19.00 | 39.00 | 18.00 | 48.00 | 53.00 | 45.00 | 20.00 |
| | 9.92 | -2.40 | 61.00 | 5.70 | 47.00 | 32.00 | 43.00 | 29.00 | 45.00 | 27.00 | 40.00 | 30.00 |
| | 65.00 | 3.10 | 37.00 | 4.60 | 46.00 | 39.00 | 45.00 | 25.00 | 52.00 | 56.00 | 41.00 | 23.00 |
| | 79.00 | 3.80 | | | 43.00 | 32.00 | 45.00 | 23.00 | 55.00 | 9.10 | 37.00 | 27.00 |
| | 73.00 | 1.20 | | | 47.00 | 42.00 | 56.00 | 15.00 | 56.00 | 13.00 | 41.00 | 17.00 |
| | 9.92 | -16.00 | | | 41.00 | 44.00 | 45.00 | 9.50 | 45.00 | 30.00 | 39.00 | 30.00 |
| | 9.92 | -15.00 | | | 39.00 | 40.00 | 42.00 | 41.00 | 42.00 | 8.40 | 35.00 | 27.00 |
| **N** | **50.00** | **50.00** | **19.00** | **19.00** | **30.00** | **30.00** | **34.00** | **34.00** | **31.00** | **31.00** | **35.00** | **35.00** |
| **AV** | **63.49** | **2.33** | **64.26** | **8.82** | **56.57** | **27.96** | **35.76** | **27.40** | **55.16** | **22.37** | **46.77** | **26.71** |
| **STD** | **27.50** | **7.43** | **13.81** | **4.26** | **11.94** | **11.69** | **20.41** | **11.85** | **8.45** | **14.08** | **9.66** | **9.82** |

N: number of observations, AV: average, STD: standard deviation.

**Table S10:** FRET efficiencies and Binding related to Fig. 4 and 5.

| **Date** | **WOX5-mV(N) PLT3-mV(N)** | | **WOX5-mV(N) PLT3-mV(N) mCherry-NLS** | | **WOX5-mV(N) PLT3-mV(N) BRAVO-mCh** | | **WOX5-mV(N) PLT3-mV(N) BES1D-mCh** | | **WOX5-mV(N) PLT3-mV(N) TPL-mCh** | |
|---|---|---|---|---|---|---|---|---|---|---|
| | **FRET E** | **Binding** | **FRET E** | **Binding** | **FRET E** | **Binding** | **FRET E** | **Binding** | **FRET E** | **Binding** |
| **16.03.2020** | 80.00 | 4.90 | | | 55.00 | 23.00 | 59.00 | 11.00 | 39.00 | 33.00 |
| | 80.00 | 6.50 | | | 50.00 | 52.00 | 53.00 | 18.00 | 39.00 | 15.00 |
| | 80.00 | 68.00 | | | 51.00 | 42.00 | 43.00 | 29.00 | 44.00 | 25.00 |
| | 80.00 | -2.30 | | | 47.00 | 34.00 | 46.00 | 19.00 | 34.00 | 34.00 |
| | 10.10 | -7.00 | | | 48.00 | 48.00 | 55.00 | 15.00 | 38.00 | 30.00 |
| | 80.00 | 8.90 | | | 46.00 | 45.00 | 61.00 | 33.50 | 40.00 | 26.00 |
| | 80.00 | 6.00 | | | 45.00 | 38.00 | 42.00 | 20.00 | 49.00 | 25.00 |

| | | | | | | | | | | |
|---|---|---|---|---|---|---|---|---|---|---|
| | 78.00 | 8.80 | | | 52.00 | 50.00 | 48.00 | 24.00 | 48.00 | 18.00 |
| | 79.00 | 4.50 | | | 49.00 | 39.00 | 45.00 | 26.00 | 43.00 | 22.00 |
| | 80.00 | 5.10 | | | 46.00 | 36.00 | 43.00 | 33.00 | 39.00 | 31.00 |
| | | | | | | | 51.00 | 35.00 | 43.00 | 34.00 |
| | | | | | | | 44.00 | 39.00 | 48.00 | 29.00 |
| | | | | | | | 44.00 | 23.00 | 44.00 | 51.00 |
| | | | | | | | 46.00 | 28.00 | 48.00 | 20.00 |
| | | | | | | | | | 41.00 | 29.00 |
| **18.12.2020** | 9.96 | -18.00 | 65.00 | 4.70 | 53.00 | 43.00 | 57.00 | 7.20 | 33.00 | 16.00 |
| | 9.96 | -11.00 | 63.00 | 7.00 | 51.00 | 45.00 | 43.00 | 15.00 | 34.00 | 26.00 |
| | 80.00 | -4.20 | 61.00 | 5.20 | 51.00 | 19.00 | 41.00 | 15.00 | 32.00 | 24.00 |
| | 9.96 | -4.70 | 51.00 | 8.80 | 43.00 | 21.00 | 49.00 | 12.00 | 37.00 | 26.00 |
| | 9.96 | -7.60 | 48.00 | 5.00 | 45.00 | 30.00 | 41.00 | 28.00 | 41.00 | 13.00 |
| | 73.00 | 2.60 | | | 48.00 | 48.00 | 45.00 | 13.00 | 33.00 | 28.00 |
| | 80.00 | 4.20 | | | 48.00 | 38.00 | 46.00 | 7.80 | 39.00 | 21.00 |
| | 54.00 | 2.20 | | | 51.00 | 25.00 | 55.00 | 6.05 | 41.00 | 14.00 |
| | 71.00 | 3.20 | | | 49.00 | 27.00 | 44.00 | 15.00 | 27.00 | 10.00 |
| | 18.00 | 3.70 | | | 45.00 | 36.00 | 39.00 | 10.00 | 35.00 | 20.00 |
| | | | | | | | 42.00 | 14.00 | | |
| | | | | | | | 50.00 | 14.00 | | |
| | | | | | | | 37.00 | 24.00 | | |
| | | | | | | | 50.00 | 12.00 | | |
| | | | | | | | 39.00 | 12.00 | | |
| **21.12.2020** | 9.95 | -15.00 | 12.00 | -2.70 | 53.00 | 42.00 | 42.00 | 18.00 | 39.00 | 17.00 |
| | 9.95 | -5.20 | 9.95 | -9.10 | 51.00 | 48.00 | 41.00 | 13.00 | 31.00 | 29.00 |
| | 10.00 | -3.60 | 10.00 | 4.30 | 46.00 | 34.00 | 56.00 | 6.90 | 43.00 | 19.00 |
| | 48.00 | -0.40 | 51.00 | 5.20 | 47.00 | 25.00 | 44.00 | 22.00 | 45.00 | 18.00 |
| | 69.00 | 2.60 | 63.00 | 1.00 | 51.00 | 12.00 | 42.00 | 17.00 | 55.00 | 9.30 |
| | 17.00 | 8.30 | | | 46.00 | 18.00 | 39.00 | 16.00 | 42.00 | 14.00 |
| | 57.00 | 5.50 | | | 51.00 | 53.00 | 36.00 | 12.00 | 36.00 | 31.00 |
| | 9.95 | -2.00 | | | 46.00 | 40.00 | 45.00 | 20.00 | 37.00 | 24.00 |
| | 9.95 | -8.30 | | | 47.00 | 40.00 | 40.00 | 20.00 | 41.00 | 15.00 |
| | 9.95 | -7.00 | | | 50.00 | 38.00 | 37.00 | 23.00 | 36.00 | 19.00 |
| | | | | | | | 41.00 | 16.00 | | |
| | | | | | | | 38.00 | 26.00 | | |
| **AV** | **46.46** | **1.62** | **43.40** | **2.94** | **48.70** | **36.30** | **45.34** | **18.74** | **39.83** | **23.29** |
| **SD** | **31.88** | **14.13** | **22.14** | **5.01** | **2.88** | **10.73** | **6.24** | **8.03** | **5.75** | **8.29** |
| **N** | **30.00** | **30.00** | **10.00** | **10.00** | **30.00** | **30.00** | **41.00** | **41.00** | **35.00** | **35.00** |

N: number of observations, AV: average, STD: standard deviation.

**Table S11:** FRET efficiencies and Binding related to Fig. S2 and 5.

| Date | BRAVO-mV(N) PLT3-mV(N) | | BRAVO-mV(N) PLT3-mV(N) mCherry-NLS | | BRAVO-mV(N) PLT3-mV(N) WOX5-mCh | | BRAVO-mV(N) PLT3-mV(N) BES1D-mCh | | BRAVO-mV(N) PLT3-mV(N) TPL-mCh | |
|---|---|---|---|---|---|---|---|---|---|---|
| | Binding | FRET E | Binding | FRET E | Binding | FRET E | Binding | FRET E | Binding | FRET E |
| 03.07.23 | 0.00 | 10.17 | 0.00 | 8.47 | 31.40 | 44.41 | 7.10 | 10.17 | 11.00 | 49.15 |
| | 0.00 | 80.03 | 2.00 | 80.03 | 40.29 | 42.37 | 0.00 | 10.17 | 16.49 | 31.53 |
| | 2.35 | 80.03 | 0.00 | 8.47 | 35.98 | 42.81 | 0.00 | 10.17 | 1.46 | 80.03 |
| | 0.00 | 10.17 | 0.00 | 80.00 | 27.89 | 46.78 | 0.00 | 79.66 | 22.00 | 13.22 |
| | 0.00 | 80.03 | 0.00 | 10.17 | 30.09 | 38.98 | 0.00 | 10.17 | 25.10 | 14.58 |
| | 0.66 | 10.17 | | | | | 2.50 | 80.03 | 8.20 | 10.17 |
| | 1.33 | 10.17 | | | | | | | 7.02 | 10.17 |
| 05.07.23 | 0.00 | 80.00 | 2.51 | 79.97 | 26.61 | 46.67 | 6.72 | 43.33 | | |
| | 0.00 | 10.00 | 0.00 | 10.00 | 26.49 | 40.67 | 11.10 | 26.67 | | |
| | 0.00 | 10.00 | 10.60 | 15.67 | 8.29 | 51.00 | 16.37 | 31.67 | | |
| | 0.00 | 79.97 | 4.10 | 60.00 | 23.92 | 33.33 | 18.80 | 38.33 | | |
| | 0.00 | 10.00 | | | 36.81 | 46.53 | 13.05 | 35.67 | | |
| | 14.20 | 10.00 | | | 43.11 | 43.00 | | | | |
| | 1.80 | 10.33 | | | 8.91 | 63.33 | | | | |
| | 0.00 | 10.00 | | | | | | | | |
| 04.12.23 | 0.00 | 10.26 | 5.34 | 14.24 | 25.03 | 36.75 | 5.25 | 80.03 | 22.54 | 31.46 |
| | 1.85 | 30.46 | 5.97 | 25.17 | 50.54 | 48.01 | 13.14 | 35.43 | 19.53 | 36.09 |
| | 4.00 | 17.22 | 3.49 | 80.03 | 42.89 | 45.03 | 15.10 | 25.17 | 34.21 | 24.83 |
| | 9.00 | 10.26 | 2.99 | 10.26 | 27.60 | 42.72 | 4.95 | 40.40 | 35.26 | 26.16 |
| | 0.00 | 10.26 | 5.00 | 80.03 | 53.65 | 51.32 | 19.30 | 35.43 | 15.13 | 42.05 |
| | 1.06 | 27.15 | | | 45.49 | 45.43 | 17.91 | 28.81 | 21.90 | 32.45 |
| | 9.30 | 11.59 | | | 12.83 | 46.69 | 15.72 | 49.34 | 31.38 | 27.15 |
| | 0.00 | 80.13 | | | 19.65 | 45.36 | 14.77 | 45.36 | 25.12 | 31.79 |
| | 8.80 | 26.82 | | | 29.73 | 43.71 | 15.30 | 37.09 | 24.00 | 37.75 |
| | 1.87 | 80.03 | | | 36.98 | 44.70 | 11.88 | 37.75 | 34.16 | 29.80 |
| 16.12.23 | 0.00 | 79.93 | 5.30 | 10.03 | 33.80 | 48.16 | 11.34 | 39.80 | 17.02 | 56.52 |
| | 0.00 | 10.03 | 4.90 | 79.97 | 20.27 | 45.48 | 9.20 | 24.75 | 22.69 | 57.19 |
| | 0.00 | 10.03 | 8.70 | 79.97 | 29.88 | 43.48 | 11.19 | 42.14 | 16.94 | 49.16 |
| | 4.40 | 10.03 | 0.00 | 10.03 | 26.93 | 39.46 | 16.09 | 42.14 | 20.00 | 48.83 |
| | 0.00 | 10.03 | 0.00 | 10.03 | 28.11 | 45.15 | 18.74 | 39.80 | 15.68 | 61.20 |
| | 0.00 | 10.03 | | | 29.75 | 45.48 | 6.64 | 29.77 | 24.97 | 36.45 |
| | 8.86 | 10.03 | | | 33.79 | 50.84 | 4.34 | 29.77 | 31.80 | 29.10 |
| | 1.50 | 80.00 | | | 20.37 | 38.13 | 10.94 | 38.46 | 17.50 | 44.48 |
| | 0.00 | 10.03 | | | 26.92 | 46.82 | 8.10 | 43.14 | 27.25 | 28.76 |
| | 7.59 | 20.74 | | | 18.77 | 45.48 | 9.75 | 29.77 | 21.80 | 39.13 |
| **AV** | 2.24 | 30.18 | 3.21 | 39.61 | 29.77 | 44.94 | 10.17 | 37.11 | 21.12 | 36.27 |
| **SD** | 3.64 | 29.78 | 3.10 | 32.79 | 10.57 | 5.13 | 5.93 | 17.45 | 8.28 | 15.98 |
| **N** | 35.00 | 35.00 | 19.00 | 19.00 | 32.00 | 32.00 | 31.00 | 31.00 | 27.00 | 27.00 |

N: number of observations, AV: average, STD: standard deviation.

**Table S12:** Additional FRET efficiencies and Binding used for Fig. 5.

| **Sample** | **WOX5-mV PLT3-mCh** | |
|---|---|---|
| **Date** | **FRET E** | **Binding** |
| **20.12.2019** | 71 | 11 |
| | 53 | 44 |
| | 57 | 34 |
| | 56 | 40 |
| | 56 | 43 |
| | 62 | 27 |
| | 55 | 48 |
| | 54 | 41 |
| | 56 | 26 |
| | 52 | 36 |
| **AV** | 57.2 | 35 |
| **SD** | 5.268776 | 10.47855 |
| **N** | 10 | 10 |

N: number of observations, AV: average, STD: standard deviation.

**Table S13:** FRET efficiencies and Binding related to Fig. 6 and 7.

| Date | BRAVO-mV | | BRAVO-mV mCherry-NLS | | BRAVO-mV PLT3-mCh | | BRAVO-mV PLT3ΔPrD-mCh | |
|---|---|---|---|---|---|---|---|---|
| | FRET E | Binding | FRET E | Binding | FRET E | Binding | FRET E | Binding |
| 23.03.2021 | 9.71 | 0.00 | | | 48 | 27 | 52 | 14 |
| | 9.71 | 0.00 | | | 51 | 19 | 49 | 23 |
| | 70.00 | 2.10 | | | 48 | 33 | 76 | 4.8 |
| | 76.00 | 3.40 | | | 58 | 8.2 | 79.9 | 5.6 |
| | 80.00 | 8.50 | | | 42 | 17 | 59 | 6.3 |
| | 10.00 | 0.00 | | | 43 | 7.7 | 79.9 | 4.7 |
| | 48.00 | 0.50 | | | 53 | 21 | 63 | 4.5 |
| | 9.71 | 0.00 | | | 45 | 35 | 43 | 7.9 |
| | 79.90 | 3.60 | | | 43 | 27 | 43 | 29 |
| | 9.71 | 0.00 | | | 56 | 22 | 43 | 44 |
| 06.04.2021 | 10.00 | -10.00 | 56.00 | 0.81 | 61.00 | 7.50 | 54.00 | 7.10 |
| | 11.00 | -3.70 | 46.00 | 1.90 | 78.00 | 3.70 | 65.00 | 2.90 |
| | 10.10 | -8.00 | 80.00 | 1.00 | 43.00 | 27.00 | 42.00 | 23.00 |
| | 15.00 | 6.20 | 67.00 | 3.30 | 47.00 | 21 | 41.00 | 18.00 |
| | 14.00 | 10.00 | 71.00 | 3.70 | 42.00 | 25.00 | 80.00 | 0.50 |
| | 10.00 | -1.50 | | | 48.00 | 24 | 9.70 | -0.80 |
| | 33.00 | 4.30 | | | 48.00 | 19.00 | 64.00 | 2.60 |
| | 80.00 | -3.60 | | | 51.00 | 19.00 | | |
| | 80.00 | 0.18 | | | 54.00 | 20.00 | | |
| | 14.00 | 5.00 | | | 46.00 | 38.00 | | |
| 21.05.2021 | 80.00 | 7.80 | 62.00 | 2.80 | 50.00 | 29.00 | 46.00 | 12 |
| | 76.00 | 5.60 | 58.00 | 8.60 | 47.00 | 13.00 | 45.00 | 21.00 |
| | 45.00 | 0.29 | 35.00 | 7.30 | 52.00 | 32.00 | 45.00 | 9.10 |
| | 69.00 | 9.00 | 61.00 | 4.80 | 49.00 | 16.00 | 34.00 | 5.60 |
| | 54.00 | 4.30 | 56.00 | 4.90 | 47.00 | 23 | 35.00 | 8.80 |
| | 36.00 | 7.10 | | | 49.00 | 29.60 | 47.00 | 6.60 |
| | 9.93 | 0.00 | | | 33.00 | 13.00 | 39.00 | 12.00 |
| | 9.93 | 0.01 | | | 47.00 | 43.00 | 37.00 | 19.00 |
| | 9.93 | 0.01 | | | 47.00 | 50.60 | 40.00 | 14.00 |
| | 9.93 | 0.01 | | | 47.00 | 15.00 | 29.00 | 14.00 |
| | | | | | | | 37.00 | 8.90 |
| AV | 35.99 | 1.70 | 59.20 | 3.91 | 49.10 | 22.84 | 49.20 | 11.72 |
| STD | 29.38 | 4.59 | 11.94 | 2.43 | 7.50 | 10.53 | 16.38 | 9.58 |
| n | 30.00 | 30.00 | 10.00 | 10.00 | 30.00 | 30.00 | 28.00 | 28.00 |

N: number of observations, AV: average, STD: standard deviation.

**Table S14:** FRET efficiency and Binding values corresponding to Figure S4.

| Date | TPL-mV | | TPL-mV mCherry-NLS | | TPL-mV PLT3-mCh | | TPL-mV PLT3ΔPrD-mCh | |
|---|---|---|---|---|---|---|---|---|
| | FRET E | Binding | FRET E | Binding | FRET E | Binding | FRET E | Binding |
| 12.04.2021 | 10.10 | 0.01 | 42.00 | 6.20 | 58.00 | 7.40 | 32.00 | 8.00 |
| | 10.10 | 0.00 | 53.00 | 3.50 | 40.00 | 16.00 | 35.00 | 7.00 |
| | 10.10 | 0.00 | 14.00 | 5.00 | 35.00 | 13.00 | 35.00 | 9 |
| | 10.00 | 0.01 | 47.00 | 6.90 | 36.00 | 8.00 | 41.00 | 7.60 |
| | 49.00 | 0.65 | 24.00 | 5.00 | 25.00 | 11.00 | 32.00 | 6.00 |
| | 55.00 | 2.20 | | | 41.00 | 16.00 | 46.00 | 12.00 |
| | 73.00 | 6.30 | | | 29.00 | 17.00 | 33.00 | 14.00 |
| | 10.10 | 0.01 | | | 30.00 | 9.50 | 50.00 | 9.7 |
| | 10.00 | 0.01 | | | 47.00 | 8.00 | 32.00 | 17.00 |
| | 30.00 | 5.10 | | | 35.00 | 12.00 | 43.00 | 11.00 |
| | | | | | 34.00 | 16.00 | 34.00 | 11.00 |
| | | | | | 25.00 | 19.00 | 36.00 | 16.00 |
| | | | | | 36.00 | 20.00 | 35.00 | 7.10 |
| | | | | | 34.00 | 18.00 | 27.00 | 6.80 |
| | | | | | 29.00 | 22.00 | 44.00 | 24.00 |
| 06.04.2021 | 10.00 | -18.00 | 74.00 | 3.10 | 43.00 | 7.70 | 35.00 | 2.60 |
| | 10.00 | -7.30 | 40.00 | 10.00 | 50.00 | 6.70 | 51.00 | 3.60 |
| | 10.00 | 5.10 | 47.00 | 7.70 | 42.00 | 12.00 | 64.00 | 2.90 |
| | 10.00 | -0.80 | 39.00 | 6.00 | 46.00 | 9.80 | 43.00 | 11.00 |
| | 10.00 | 8.60 | 48.00 | 11.00 | 42.00 | 16.00 | 59.00 | 3.50 |
| | 11.00 | 3.40 | | | 46.00 | 11.00 | 26.00 | 4.60 |
| | 15.00 | 1.80 | | | 48.00 | 15.00 | 70.00 | 5.80 |
| | 10.00 | -2.00 | | | 44.00 | 17.00 | 47.00 | 13.00 |
| | 27.00 | 0.86 | | | 47.00 | 19.00 | 47.00 | 12.00 |
| | 10.00 | 6.80 | | | 29.00 | 18.00 | 80.00 | -0.48 |
| **AV** | 19.52 | 0.64 | 42.80 | 6.44 | 38.84 | 13.80 | 43.08 | 8.99 |
| **STD** | 17.91 | 5.51 | 15.32 | 2.44 | 8.31 | 4.42 | 13.19 | 5.26 |
| **n** | 20.00 | 20.00 | 10.00 | 10.00 | 25.00 | 25.00 | 25.00 | 25.00 |

N: number of observations, AV: average, STD: standard deviation.

**Table S15:** FRET efficiency and Binding values corresponding to Fig. S4.

| Date | BES1-mV | | BES1-mV mCherry-NLS | | BES1-mV PLT3-mCh | | BES1-mV PLT3ΔPrD-mCh | |
|---|---|---|---|---|---|---|---|---|
| | FRET E | Binding | FRET E | Binding | FRET E | Binding | FRET E | Binding |
| 21.05.2021 | 9.91 | -10.00 | 80.00 | 20.00 | 68.00 | 16.00 | 34.00 | 13.00 |
| | 10.00 | -1.30 | 48.00 | 2.60 | 48.00 | 7.70 | 44.00 | 7.10 |
| | 80.00 | 0.95 | 46.00 | 5.70 | 45.00 | 9.90 | 54.00 | 8.00 |
| | 39.00 | 0.78 | 45.00 | 12.00 | | | | |
| | 9.91 | -2.40 | 50.00 | 7.60 | | | | |
| | 54.00 | 3.40 | | | | | | |
| | 18.00 | 4.40 | | | | | | |
| | 9.91 | -13.00 | | | | | | |
| | 9.91 | -12.00 | | | | | | |
| | 9.91 | -5.60 | | | | | | |
| 27.04.2021 | 80.00 | 3.40 | 39.00 | 0.73 | 76.00 | 6.50 | 57.00 | 4.40 |
| | 80.00 | 0.00 | 60.00 | 5.70 | 59.00 | 8.90 | 54.00 | 6.30 |
| | 10.10 | 0.00 | 80.10 | 4.30 | 48.00 | 19.00 | 57.00 | 7.10 |
| | 10.10 | 0.00 | 80.00 | 1.00 | 43.00 | 23.00 | 45.00 | 18.00 |
| | 59.00 | 4.40 | 65.00 | 3.20 | 51.00 | 24.00 | 47.00 | 18.00 |
| | 67.00 | 4.40 | | | 45.00 | 35.00 | 13.00 | 2.20 |
| | 70.00 | 9.20 | | | 66.00 | 8.10 | 76.00 | 6.00 |
| | 55.00 | 9.70 | | | 55.00 | 6.10 | 56.00 | 3.50 |
| | 80.00 | 2.30 | | | 49.00 | 9.90 | 67.00 | 6.70 |
| | 80.00 | 2.00 | | | 46.00 | 22.00 | 51.00 | 8.3 |
| 28.05.2021 | 80.00 | -3.50 | 10.00 | -12.00 | 56.00 | 9.90 | | |
| | 80.00 | 0.67 | 77.00 | 3.2 | 41.00 | 18.00 | | |
| | 10.00 | -12.00 | 63.00 | 7.00 | 45.00 | 20.00 | | |
| | 80.00 | 7.50 | 49.00 | 7.00 | 47.00 | 26.00 | | |
| | 80.00 | -1.40 | 48.00 | 5.60 | 65.00 | 7.30 | | |
| | 10.00 | -12.00 | | | 42.00 | 23.00 | | |
| | 80.00 | 1.40 | | | 42.00 | 27.00 | | |
| | 80.00 | 6.60 | | | 45.00 | 28.00 | | |
| | 57.00 | 7.20 | | | 48.00 | 19.00 | | |
| | 64.00 | 5.10 | | | 46.00 | 14.00 | | |
| **AV** | 48.76 | 0.01 | 56.01 | 4.91 | 51.13 | 16.88 | 50.38 | 8.35 |
| **STD** | 30.59 | 6.34 | 18.55 | 6.46 | 9.34 | 8.09 | 14.74 | 4.81 |
| **n** | 30.00 | 30.00 | 15.00 | 15.00 | 23.00 | 23.00 | 13.00 | 13.00 |

N: number of observations, AV: average, STD: standard deviation.

**Table S16:** Ratio of periclinal cell divisions in the QC related to Fig. 7.

| Treatment | Genotype | Periclinal cell division planes in the QC [%] | Number of analysed roots |
|---|---|---|---|
| DMSO | *Col-0* | 27 | 44 |
| | *plt3-1* | 73 | 45 |
| | *plt3-1, pWOX5:GR-PLT3-mT2* | 83 | 41 |
| | *plt3-1, pWOX5:GR-PLT3dPrD-mT2* | 94 | 36 |
| DEX | *Col-0* | 28 | 39 |
| | *plt3-1* | 87 | 46 |
| | *plt3-1, pWOX5:GR-PLT3-mT2* | 67 | 45 |
| | *plt3-1, pWOX5:GR-PLT3dPrD-mT2* | 100 | 36 |
